# A tractable bottom-up model of the yeast polarity genotype-phenotype map for evolutionary relevant predictions

**DOI:** 10.1101/2020.11.09.374363

**Authors:** Werner Karl-Gustav Daalman, Els Sweep, Liedewij Laan

## Abstract

Accurate phenotype prediction based on genetic information has numerous societal applications, such as crop design or cellular factories. Epistasis, when biological components interact, complicates modelling phenotypes from genotypes. Here we show an approach to mitigate this complication for polarity establishment in budding yeast, where mechanistic information is abundant. We coarse-grain molecular interactions into a so-called mesotype, which we combine with gene expression noise into a physical cell cycle model. Firstly, we show with computer simulations that the mesotype allows validation of the most current biochemical polarity models by quantitatively matching doubling times. Secondly, the mesotype elucidates epistasis emergence as exemplified by evaluating the predicted mutational effect of key polarity protein Bem1p when combined with known interactors or under different growth conditions. This example also illustrates how unlikely evolutionary trajectories can become more accessible. The tractability of our biophysically justifiable approach inspires a road-map towards bottom-up phenotype modelling beyond statistical inferences.

## Introduction

Many fields, such as personalized medicine (Li, 2011), agriculture (Cobb et al., 2013), chemical production (Oud et al., 2012) and forensics (Kayser and Schneider, 2009), will greatly benefit from better understanding of how traits connect to genes, the so-called genotype-phenotype (GP-) map. However, resolving this connection is generally not straightforward even for known heritable traits (Eichler et al., 2010). For example, multiple genes can contribute to a single trait (polygenic inheritance) while multiple traits can emerge from a single gene (pleiotropy) (Wagner and Zhang, 2011). Frequently, mutational effects have been shown to be non-additive in (model) species as humans (Cordell, 2009; Zuk et al., 2012), *Escherichia coli* (Wünsche et al., 2017) and *Saccharomyces cerevisiae* (budding yeast) (van Leeuwen et al., 2016). This phenomenon is known as epistasis. Theoretically, epistasis surfaces easily based on metabolic network analysis (Kryazhimskiy, 2021), and some molecular origins are known (Lehner, 2011). Although sometimes inconsequential for fitness evolution (Kryazhimskiy et al., 2014), epistasis complicates the predictions of phenotype and consequently gene evolution (Papp et al., 2011; Miton and Tokuriki, 2016). Therefore, epistasis predictions pose an important challenge for GP-map models.

In order to unravel the complication epistasis imposes, intermediate levels in the GP-map are commonly employed (Blanco-Gómez et al., 2016). An intermediate level is any quantity more complex than individual proteins, but less complex than the phenotype level. This level addition is meant to produce a more modular and hence more tractable GP-map (Figure 1A). Ideally, a level that clarifies trait generation also elucidates the handles for evolution, the reverse path in the GP-map. While phenomenological or statistical level formulations may suffice for predictions, they are not designed to provide insight into the GP-map. We therefore formulate a biophysically sound intermediate level from the bottom-up, which we refer to as the “mesotype”.

**Figure 1.**
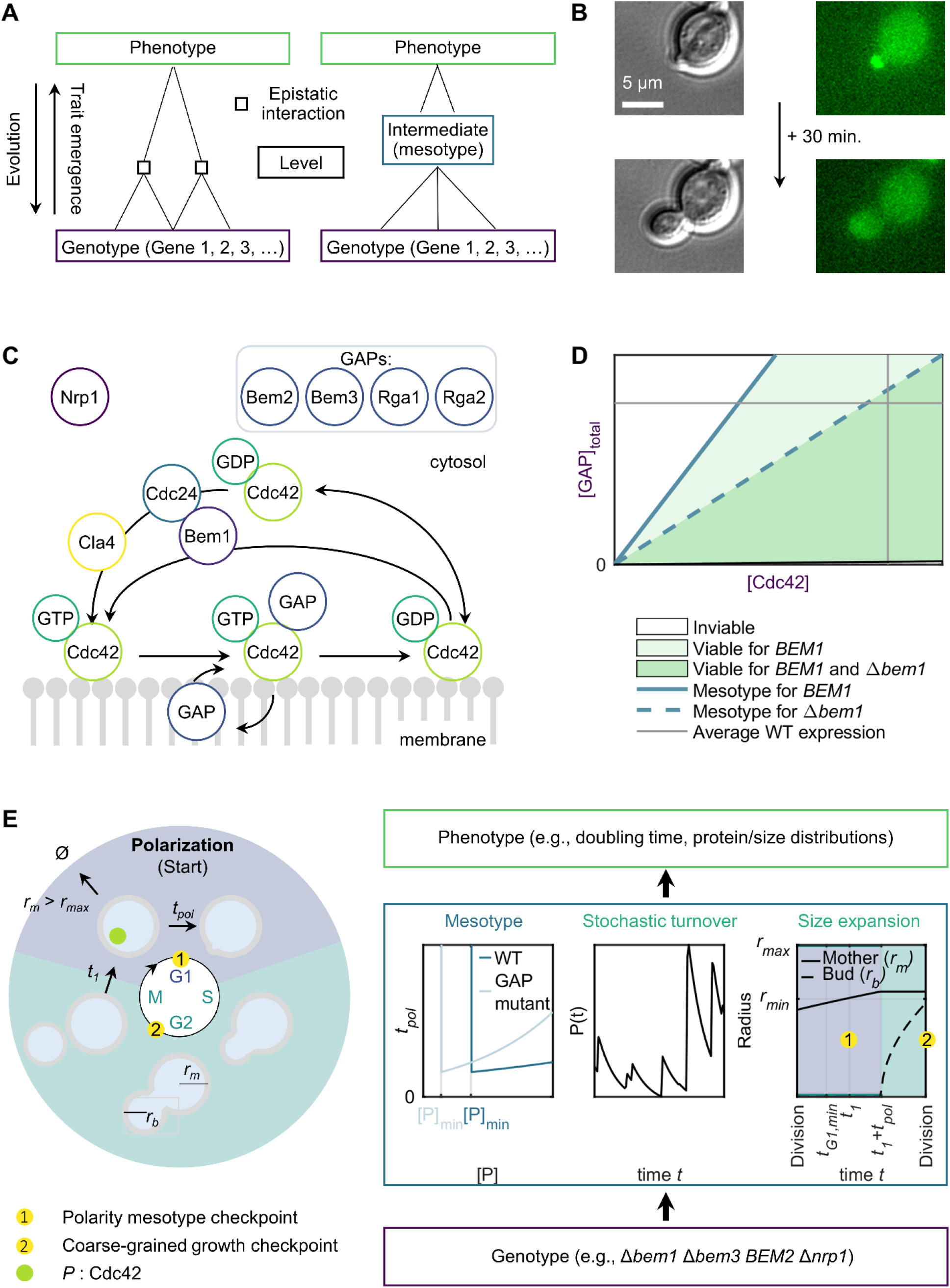
Decomposition of the genotype-phenotype map with the mesotype as an intermediate, showcased in yeast polarity. (A) General conceptualization of a genotype-phenotype (GP-) map containing epistatic interactions. Introduction of an intermediate level (right) ideally simplifies the connections between genes to phenotype, improving tractability for trait generation (upwards trajectory) and evolution (coupling downwards from the phenotype). (B) Visualization of yeast polarity as the function template for decomposing a GP-map. Brightfield (left) and widefield fluorescence (right) example images of a polarizing budding yeast cell (yLL129), scale bar 5 µm. Key for polarity is clustering to one zone on the membrane of active Cdc42p, of which a binding partner (Spa2p) is fluorescently labelled in the images. This zone marks the site of polarized growth. (C) Schematic model overview of core polarity protein network (proteins not to scale). A positive feedback for (active) Cdc42p-GTP is mediated by the Bem1p-Cdc24p complex, and to lesser extent by Cla4p. Nrp1p represents a mechanistic unknown. (D) Phase diagram summarizing the GP-map, depicting the phenotype viability (green) as function of genotype (purple) through Cdc42p (active and inactive) and GAP concentration in the cell with or without Bem1p. The GP-map contains epistatic interactions, for example the same increase in GAP concentration can yield inviability in the Δ*bem1* background but not in the *BEM1* background. An intermediate, the ‘mesotype’, can be identified here as the limiting Cdc42p concentration (blue). (E) Implementation of the mesotype into a physical cell cycle model, to tractably decompose the polarity GP-map. Starting from G1, every cell aims to divide once the polarity mesotype checkpoint has passed. This implies mother radius *r_m_* exceeding minimum radius *r_min_*, time thus far in G1 *t_1_* exceeding minimum G1 time *t_G1,min_*, and a Cdc42 concentration (abbreviated as [*P*]) exceeding mesotype [*P*]*_min_*. If maximum size *r_max_* is exceeded in the process, the cell dies. If the mesotype checkpoint is passed, the mother continues to grow isotropically for polarization time *t_pol_*, which is at least *t_pol,min_*. Then, only the bud grows until the second, coarse-grained growth checkpoint, which involves bud size must be a certain fraction of mother size before division. Throughout all phases, Cdc42p is subject to stochastic protein production and deterministic degradation. See also Table S1.

To test our mesotype approach, we model polarity establishment in *S. cerevisiae*, the cell cycle step where budding yeast breaks its spherical symmetry to direct bud growth (see Figure 1B). The challenge is reproducing the ample epistasis exhibited in the polarity network in e.g., doubling times (Laan et al., 2015). Conveniently, the molecular interaction network for polarity establishment (simplified in Figure 1C) was recently modelled in broad agreement with literature (Brauns et al., 2020). In short, polarity is essential for cell proliferation and relies on the small GTPase Cdc42p (Adams et al., 1990). To cluster the active form of Cdc42p, bound to a GTP molecule, on one plasma membrane patch, mutual recruitment of Cdc42p and Bem1p complexes containing Cdc24p occurs (Goryachev and Pokhilko, 2008; Kozubowski et al., 2008; Klünder et al., 2013; Witte et al., 2017). Cdc24p activates Cdc42p locally (Woods et al., 2015), while Cdc42p is inactivated by GTPase activating proteins (GAPs), namely Bem2p, Bem3p (Zheng et al., 1994), Rga1p (Stevenson et al., 1995) and Rga2p (Smith et al., 2002). In absence of Bem1p, transport presumably occurs by actin (Freisinger et al., 2013) and/or Cla4p (Tiedje et al., 2008). Clusters of active Cdc42p arise by locally saturating the GAPs (Brauns et al., 2020). In the Δ*bem1* background, polarity establishment benefits from deleting the gene *NRP1*, possibly by influencing cell cycle timing (Laan et al., 2015). However, Nrp1p’s molecular mechanism is currently unknown.

Based on these molecular details and associated rigorous analysis of reaction-diffusion equations, we construct the genotype-phenotype map for yeast cell polarity by coarse-graining. One particular result in (Brauns et al., 2020) motivates simplifying polarity success to certain GAP/Cdc42p dosage ratios (Figure 1D). More specifically, a minimum Cdc42p concentration exists below which polarization is not possible. This simple rule on protein dosage defines the mesotype in this context and facilitates the understanding of epistasis emergence while sharply reducing computational costs.

To complete our GP-map model, the mesotype is incorporated into a physical cell cycle model, which is further composed of simple volume growth and stochastic protein production (Figure 1E), to reproduce phenotypes as epistasis. In this paper we assess the quality of our predictions by comparing these to documented interactions, validating the underlying molecular polarity model quantitatively. Our tractable approach also illustrates how epistasis and therefore feasible evolutionary trajectories depend on growth conditions. Finally, we show our framework relies on biofunctional (and ideally mechanistic information) of the key proteins to yield informative predictions, which delineates the appropriate conditions for applying our method.

## Results

### Cell cycle model design with the mesotype as level between genotype and phenotype

We modelled the yeast cell cycle as a process involving three modules, namely (I) polarity mesotype, (II) stochastic protein turnover and (III) cell size expansion (Figure 1E). We integrated these modules into a simulated population of cells, with properties size and protein content. In a Gillespie-style algorithm, the properties are updated per cell (Figure 2A) to yield population dynamics (Figure 2B). We sketch the essence of each module, and further details are found in the Materials and Methods. Moreover, a summary of model parameter values is given in Table S1.

**Figure 2.**
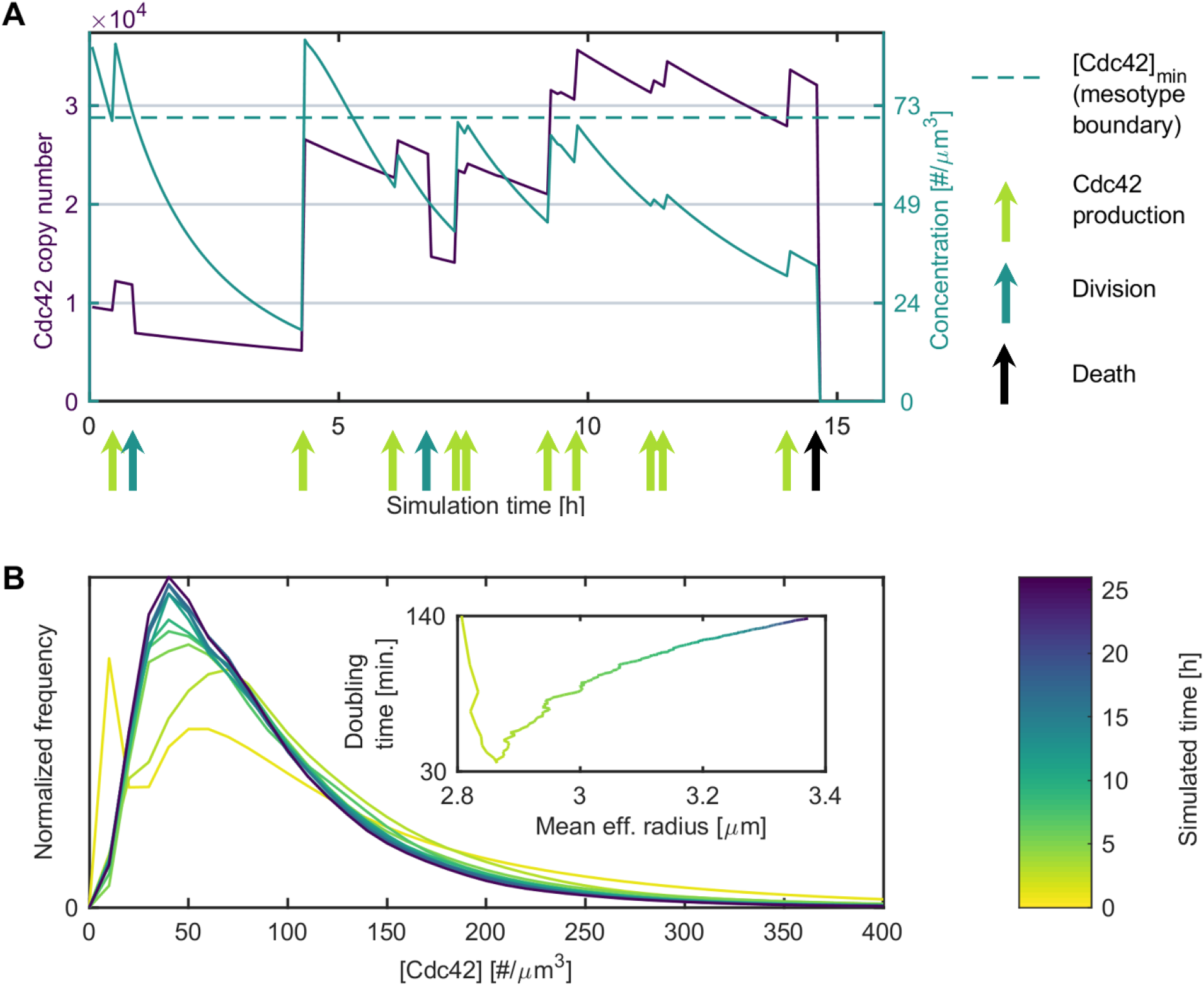
Verification of our physical cell cycle model. (A) Example trace of the Cdc42 copy number (purple) and concentration (emerald) of a single cell, which are subject to protein production and degradation (and dilution for concentrations). The cell must exceed the mesotype threshold ([Cdc42p]*_min_*) before division can take place. When this is delayed for too long, the cell expands beyond the predefined maximal size *r_max_* and dies (after almost 15h). (B) Convergence of Cdc42 copy number distribution during simulations. Simulated time since ancestor is approximate as birth times of the cells in the starting population are distributed across an 83 min. bandwidth. The inset shows how the estimates of the population doubling time and the average effective cell size equilibrate as a function of time. See also Figure S1 concerning the doubling time convergence, and Figure S3 concerning measured stochasticity of Cdc42p expression.

For module I, we reduced the biochemical network for polarity to the mesotype of a minimal [Cdc42p] threshold (dotted line Figure 2A). GAP mutations shift this threshold downward, which in part constitutes the polarity mesotype checkpoint (see module III). Upon checkpoint passage, cell expansion continues for a time *t_pol_*, which increases exponentially with the excess [Cdc42p]. This time period simplifies the functional dependency uncovered with the aforementioned analysis of the underlying reaction-diffusion equations (Brauns et al., 2020).

For module II, we only consider Cdc42p to induce cell-to-cell variability, as GAP dosage population noise (Chong et al., 2015) is much smaller than the noise of Cdc42p (coefficient of variation 0.83, this study). As mRNA of Cdc42 lives much shorter than the protein (Christiano et al., 2014; Grigull et al., 2004), we assume bursty Cdc42p production (Figure 2A). By contrast, we assume deterministic Cdc42p degradation due to its high abundance (Kulak et al., 2014).

For module III, we assume two stages of constant outer membrane area growth in G1 and S/G2/M phase respectively. In the first stage, spherical mother cells of radius *r_m_(t)* grow until the polarity mesotype checkpoint is passed and polarization is completed, or until cell death when *r_m_(t)* exceeds maximum radius *r_max_* (black arrow Figure 2A). The polarity checkpoint entails next to the [Cdc42p]*_min_* of module I, exceeding a minimal radius *r_min_*, and a minimal time since last division *t_G1,min_*,. The last two criteria represent e.g., cell size dependent control of Cln3p nuclear arrival by Ydj1p (Vergés et al., 2007). The *nrp1* deletion is phenomenologically incorporated by reducing this minimal time as suggested by Laan et al., 2015). In the next stage, a bud with radius *r_b_(t)* grows to 70% of the mother volume (the second checkpoint). Then, mother and bud restart the cell cycle independently. Resulting cell expansion rates are in decent agreement with literature (Ferrezuelo et al., 2012; Goranov et al., 2009), see Materials and Methods: size expansion.

Finally, we validated the convergence of the phenotypes dosage distribution, size and doubling times (Figure 2B). In particular, the size and doubling time change 0.6% and 0.9% respectively across the last 200 minutes, well within the typical experimental error of (Laan et al., 2015). Confirmation of doubling time convergence is also depicted in Figure S1.

### The mesotype enables detailed validation of the underlying biochemical model and bottom-up interpretation of phenotype emergence

Our physical cell cycle model relies on the validity of the underlying biochemical network model, which was coarse-grained to the mesotype. While the biochemical network model was tested in (Brauns et al., 2020), our cell cycle model allows otherwise unfeasible quantitative phenotype descriptions, improving the validation precision of the biochemical network model.

To this end, we considered 20 previously documented experimental values from (Laan et al., 2015). These constitute doubling times, whose reciprocals are defined as fitness, and cell cycle times. The three most prominent, non-trivial phenotypes are (i) strong epistasis in growth rates between GAP mutants only in the Δ*bem1* background, (ii) strong epistasis between *BEM1* and *NRP1*, and (iii) non-monotonous optimization of G1 times for (reconstructed) experimentally evolved mutants starting from Δ*bem1*. For the latter phenotype, the acceleration of G1 speed of the Δ*bem1* cells, despite their poor fitness, compared to WT cells is particularly noteworthy.

To model the mutants in our target observable set, we required five fitting parameters. Firstly, the *BEM1* and Δ*bem1* have two different values for the mesotype. Secondly, because the mesotype scales with GAP concentration (Figure 1D), any combination of *bem2* and *bem3* null mutations is described by two additional parameters. Lastly, the *nrp1* mutant is phenomenologically incorporated through a (25%) decrease in *t_G1,min_*.

The first phenotype of interest, GAP epistasis, is quantitatively well described (Figure 3A, left). This description is robust to several model assumption modifications (Figure S2). Particularly in presence of *NRP1*, fitness values were modelled in accurate accordance with experiments of (Laan et al., 2015). As expected, less cells produce sufficient Cdc42p for high mesotypes, leading to increased cell death and lower fitness (Figure 3B). Therefore, we see a diffuse viability threshold despite a sharp mesotype, as in previous experiments with inducible Cdc42p (Brauns et al., 2020).

**Figure 3.**
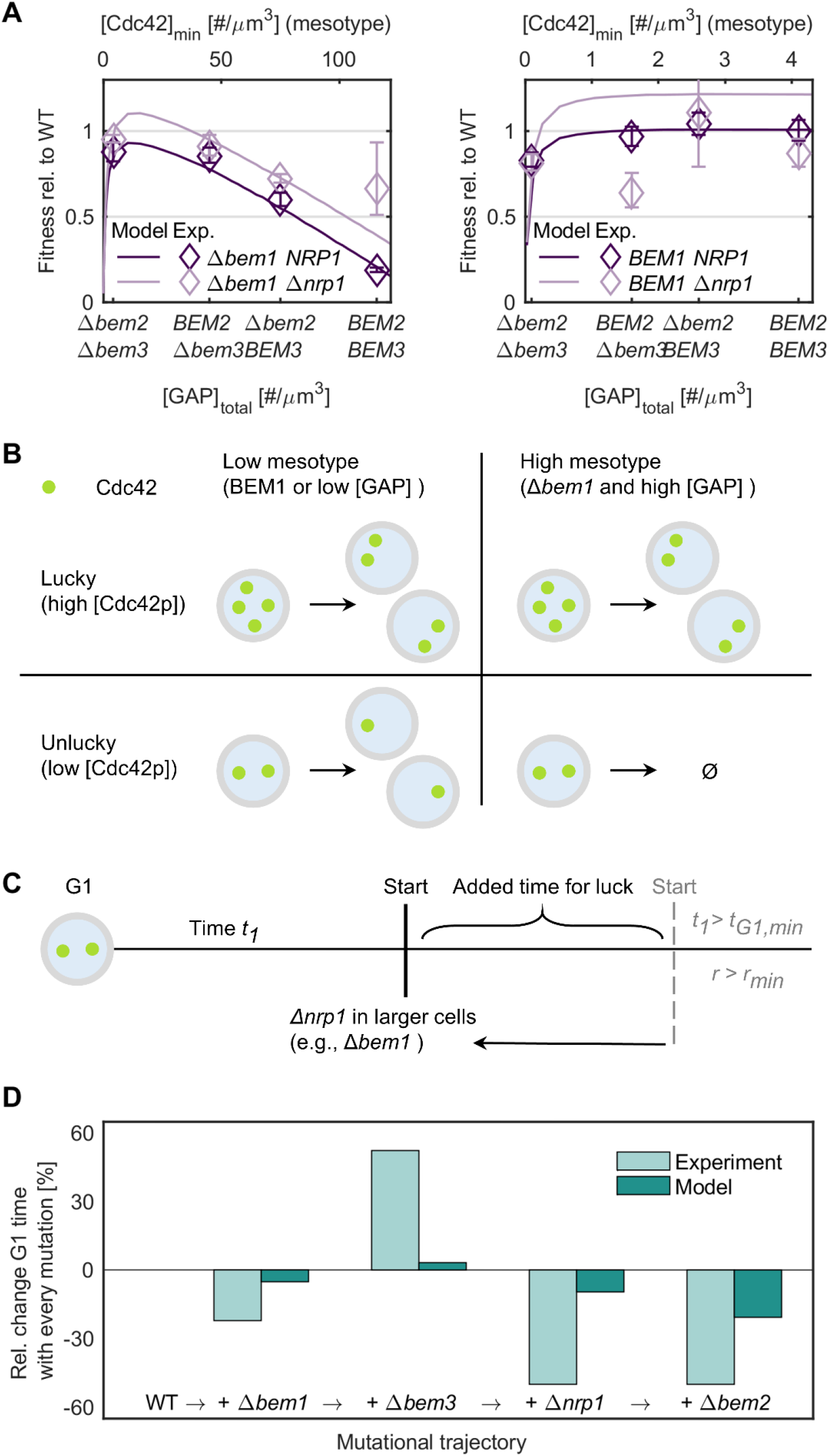
Our physical cell cycle model including the mesotype permits detailed and tractable validation of the underlying biochemical polarity model. (A) Experimental fitness values relative to WT (phenotypes) for 16 different polarity genotypes (Laan et al., 2015) denoted by diamonds, which are fitted by our physical cell cycle model as depicted by the dark and light lines for Δ*nrp1* and *NRP1* background respectively. GAP genotypes can be linearly linked to the minimum Cdc42 concentration to polarize, the mesotype, as displayed on the top horizontal axis. Error of the *BEM2 NRP1* was not available and conservatively guessed. See also Figure S2. (B) Graphical representation of the effects of GAP and *bem1* deletions from the mesotype perspective. The stochasticity of exceeding the minimal [Cdc42p], the mesotype, only becomes apparent in absence of Bem1p and with high GAP concentrations. (C) Graphical representation of the effects of the *nrp1* deletion from the mesotype perspective. When cells are less fit and become larger, the minimal G1 time reduction associated with the *nrp1* deletion becomes most apparent. Otherwise, Start is more frequently set by the minimum size criterion. The added time for lucky overproduction of Cdc42p is beneficial for the backgrounds that suffer from Cdc42p underproduction, like the Δ*bem1*. (D) The detailed phenotype of minimum G1 time as displayed by WT and four polarity mutants, comprising a single evolutionary trajectory. Experimental values are from (Laan et al., 2015) in light emerald (defined there as tie to first polarity spot), model values are in dark emerald (defined here as the time in G1 until both the size and time criteria are met). Both cases are normalized to their respective WT values, such that each column denoted the relative change in G1 time compared to the previous step in the trajectory. See also Table S2.

The four fitted mesotype thresholds are consistent with a Δ*bem3* effect that is twice as large as for Δ*bem2*. Given the GAP abundancies (Kulak et al., 2014), this sets the *in vivo* Bem3p effective GAP activity to be almost four times that of Bem2p. This robust result (Figure S2) suggests that the GAPs are functionally more similar *in vivo* than measured *in vitro,* where Bem2p activity was negligible compared to Bemp3 activity. (Zheng et al., 1994).

If we consider the second phenotype of interest, *BEM1*-*NRP1* epistasis, we focus attention to the Δ*nrp1* background. There, fitness values (Figure 3A, right) are not always well fitted (5/8 correct within experimental error), although these mutants have relatively large experimental uncertainties. Nevertheless, the strongest feature, the *BEM1*-*NRP1* epistasis, is at least qualitatively described. This description confirms the intuition offered by the mesotype. If the *nrp1* deletion reduces the mandatory G1 waiting time *t_G1,min_*, Δ*bem1* cells have more chance to exploit temporary Cdc42p overproduction before excessive dilution, thereby improving fitness (Figure 3C). Quantitatively, we explain one-third of the epistasis following the definition of (da Silva et al., 2010). An alternative G1 time formulation cannot strongly improve this result (see Materials and Methods, G1 timing). In the next section, we confirm that incorporating mutants phenomenologically rather than based on molecular information limits the accuracy of phenotype description.

Lastly, we turned to the third phenotype of interest concerning non-monotonous G1-time shifts during adaptation. These shifts are an example of the more detailed traits that can be modelled. To mimic the poorer content of synthetic medium in (Laan et al., 2015), we performed the simulations with half the normal membrane area rates. The observed trends in G1 times along the evolutionary trajectory from WT to the fully evolved mutant in that paper were qualitatively matched, including the unusual G1 time decrease for the Δ*bem1* (Figure 3D). The mesotype clarifies this subtle phenotype. The Δ*bem1* cells are relatively larger and less limited by the minimum size criterion *r_min_*. This eliminates a potential waiting step in G1 for this background, which may allow the cells to pass G1 faster if these have lucky Cdc42p overproduction. Other phases are on average extended by the relatively high minimum Cdc42 concentration threshold.

In summary, the three phenotype examples illustrate two main points. Firstly, comparison of experimental data with simulation of our physical cell cycle model further confirms the current biochemical model view of polarity. Secondly, the mesotype framework exhibits the tractability needed for understanding phenotype emergence, as demonstrated with several examples.

### The mesotype generates model predictions for genetic interactions

#### Poorer medium quality reduces fitness differences

After establishing the value of the mesotype in trait generation, we turn to the reverse transition in the GP-map. This transition is embodied by evolution (Figure 1A), since phenotypes such as fitness determine the selective pressure to shape genotypes. Environmental factors are important for growth, with even subtle changes noticeable under highly controlled laboratory settings (Atolia et al., 2020). Given that historical evolution has occurred in the wild, where conditions are expected to be much more variable and more often difficult than not, adaptive trajectories can be very different than for laboratory evolution. For example, step-wise evolution of Bem1p has already been hypothesized (Brauns et al., 2020) given fitness in lab conditions. Yet, more difficult conditions, such as slower growth and concordant ease of exceeding the mesotype, suggests that the evolution of cells lacking Bem1p may be more likely than initially anticipated.

To further substantiate our mesotype intuition on condition-dependence for Bem1p evolution, we first simulate the effect of changes in growth media quality through a change in membrane area growth rates. We considered a roughly three-fold area growth rate range that caused WT fitness to span between 0.5 and 1 (normalized to maximum growth). We further assume Cdc42p expression remains the same across media, which is at least true upon switching from dextrose to ethanol, an inferior carbon source (Smith and Kruglyak, 2008).

As a result of our model simulations, Figure 4A displays the fitness ratio between the Δ*bem1* and *BEM1* background, as a function of GAP concentration and medium quality. Intuitively, we expected Δ*bem1* cells to benefit greatly from less Cdc42p dilution and the extra time to exceed the mesotype threshold. We indeed observe the trend of smaller fitness differences for decreasing GAP concentrations and decreasing medium quality in our simulations. This observation suggests poorer media acts as a fitness equalizer, facilitating the evolution of Bem1p.

**Figure 4.**
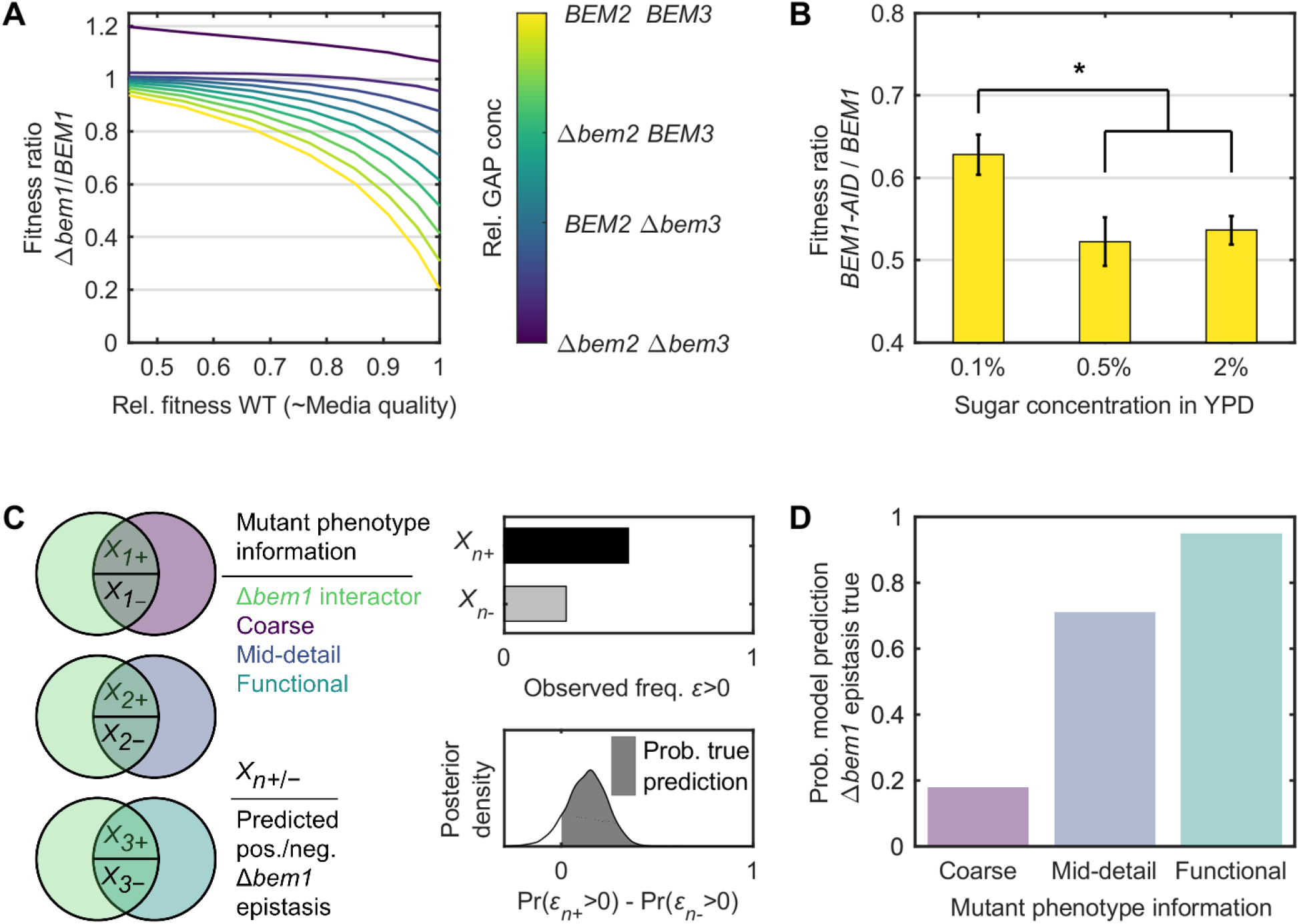
The mesotype allows evolutionary relevant predictions on the effect of medium quality and epistasis. (A) Simulated fitness differences between *BEM1* and Δ*bem1* backgrounds as a function of medium quality, which is integrated in the cell cycle model through varying cell membrane area growth rates. The medium quality translates to an associated WT fitness, which sets the horizontal axis. Different GAP concentrations are denoted by different colors. Generally, poorer medium reduces differences in fitness and genetic interactions between GAPs when comparing the *BEM1* and Δ*bem1* backgrounds. (B) Experimental assay comparing the effect of the *bem1* deletion in rich and poor medium. Bars denote the fitness is plotted of the effective Δ*bem1* background (*BEM1-AID* with added auxin) relative to the *BEM1* background (all with *BEM2* and *BEM3*), for three medium conditions. From left to right, quality of the YPD medium increases, with increasing dextrose concentration as indicated on the axis. Asterisks denote significant differences. (C) Workflow for model prediction on epistasis *ε* of various mutants with Δ*bem1*. Mutants are divided into three sets by the intersection of Bem1 interactors with mutants of various phenotype specificity (and hence varying model implementation). Each set consists of two subsets, depending on the subsequent model prediction on epistasis sign. For each subset, the beta posterior density of the observed positive epistasis fraction can be constructed (from a binomial likelihood and uniform prior). The model prediction given the information per set is that + subset has more positive epistasis than - subset. The probability of a true prediction is then defined as the area below the posterior density of the difference in positive epistasis abundance in both subsets. See also Table S3. (D) Probability bars reflecting Bayes factors for the model hypothesis; the ratio between the odds that the model prediction on positive epistasis abundance differences in subsets is true and false.

To experimentally test our hypothesized environmental effect, Figure 4B demonstrates the effect of poorer medium on fitness with and without Bem1p. In this case, we modulated medium quality through sugar content (from 2% to 0.1%). To avoid lengthy exposure of the Δ*bem1* background to high selective pressure, this genotype is mimicked by auxin inducible degradation (Nishimura et al., 2009) of Bem1p. As it is difficult to exactly integrate the media conditions into the simulations, the match cannot be expected to be quantitative. However, the nullifying effect of poor media on the fitness differences between the Δ*bem1* and *BEM1* backgrounds is visible. The relative fitness compared to WT for the effective Δ*bem1* background in the ‘poorest’ media (0.1% dextrose) is significantly better compared to the ‘richest’ and intermediate medium conditions at 2% and 0.5% dextrose (one-sided Welch’s t-test, p-value 1.3·10^-3^ and 3.3·10^-3^ respectively), also when considering the Holm-Bonferroni correction (Holm, 1979). Thus, the mesotype framework can provide a new, intuitive explanation of otherwise non-trivial environmental interactions that are relevant for evolution.

#### The polarity mesotype predictions on epistasis become useful when functions of mutated genes are known

While we showed that our cell cycle model produced accurate epistasis predictions, the mechanistic information we used is not always available. We therefore determined the required mutant information content for qualitative epistasis predictions with modules where no mechanistic information is assumed. For this purpose, we considered high-throughput data on *BEM1* interactors, assuming many are documented given the strong *bem1* phenotype. Intersecting *BEM1* interactors and mutants with various phenotype specificities, i.e. information content, yields three sets. (Figure 4C). From low to high information content, the coarse phenotype set covers (competitive/fermentative) fitness mutants, the mid-detail set G1 mutants (in size/speed), and the functional set covers proteasomal, phospholipid or ribosomal mutants. Our model subdivides each set by sign of the predicted epistasis, positive (*X_n_*_+_) or negative (*X_n_*_-_). These predictions can be intuitively constructed as explained in the next paragraphs or derived from simulations (Table S3). We then determined for which set, and thus information content, the observed epistasis sign prevalence qualitatively matched predictions (see Materials and Methods, quantification and statistical analysis).

Firstly, the coarse phenotype mutants are incorporated in the membrane area growth rate *C_2_*. Similar to the media effect in Figure 4A, smaller rates mitigate the Δ*bem1* effect. The model prediction is hence that epistasis with Δ*bem1* for deleterious mutants (*ε_1-_*), which slow down growth, is more often positive than the epistasis for beneficial mutants (*ε_1+_*), We require a Bayesian odds ratio Pr(*ε_1+_* > 0)/Pr(*ε_1-_* > 0) > 10 (Kass and Raftery, 1995) or equivalently, a probability that our epistasis statement is true of > 90 %. However, in the literature data only 14% of the deleterious mutants have positive epistasis with Δ*bem1*, compared to 19% of the beneficial mutants. Consequently, the data shows no support for our statement on epistasis prevalence given coarse mutant information, with the 18% chance (Figure 4D) worse than a coin-flip (50%).

Secondly, the mid-detail mutants are incorporated by changing G1 waiting time *t_G1,min_* (for G1 speed mutants) and additionally minimal radius before Start *r_min_* (for G1 size mutants). As was the case for to Nrp1p (Figure 3C), mutations which shorten Start disproportionally benefit the Δ*bem1* background. Therefore, the model prediction is that mutants fast or small in G1 have more positive epistasis with Δ*bem1* than mutants that are slow or large in G1. While positive epistasis is more abundant for fast/small in G1 mutants (27% against 20%), the experimental evidence is not yet compelling, with 71% chance that this model claim is true.

Lastly, the functional mutant set is incorporated by increasing the Cdc42p half-life for proteasomal mutants, decreasing membrane growth rate *C_2_* for phospholipid mutants and decreasing average Cdc42p burst size for ribosomal mutants. The proteasomal and phospholipid mutants mitigate the problematic lack of Cdc42p in the Δ*bem1* cells. These two mutant types should therefore exhibit more positive epistasis than the ribosomal mutants, which lowers [Cdc42p]. From the literature data, 29% of the proteasomal and phospholipid mutants have positive epistasis, much more than the 11% of the ribosomal mutants. There is hence strong positive evidence for our epistasis prevalence statement, which is true with around 95% certainty. This shows we minimally require functional information on mutants for meaningful epistasis predictions, once we have a core where mechanical information is known.

## Discussion

Epistasis forms a general hurdle for reverse-engineering the GP-map, complicating modelling of protein networks and in turn limiting the predictability of phenotypes and evolution. To alleviate the GP-map complexity, we tested the mesotype as an intermediate level between genotype and phenotype. This level emerges from coarse-graining the biophysics of the underlying biochemical network, which in our case corresponded to budding yeast polarity (Figure 1). We employed the mesotype, which in our context is the minimum [Cdc42p] to polarize, in a tractable cell cycle model with simple volume growth and stochastic protein production, making bottom-up reconstruction of phenotypes and epistasis feasible and insightful.

Firstly, we validated the polarity network model to unprecedented detail by our cell cycle model. Good quantitative agreement with literature was found for 13 out of 16 polarity mutant doubling times, and qualitative agreement for documented unintuitive G1 times (Figure 3). Phenomenological linkage of minimum G1 time to Nrp1p, a protein normally not included in polarity models, also provided fruitful model predictions. Thus, our cell cycle model also provides direction to deciphering the remaining mechanistic unknowns in the polarity network.

Secondly, we showed how the mesotype elucidates epistasis. We hypothesize that lower quality growth medium extends protein production time, alleviating minimum protein concentration thresholds. We verified this for yeast polarity, where the [Cdc42p] mesotype threshold proves less problematic for the Δ*bem1* background in poorer media in simulations and experiments (Figure 4A and 4B). The Δ*bem1* case fits the more general picture that haploinsufficiency in YPD is typically lifted in poorer medium (Deutschbauer et al., 2005). By the same token, evolvability of (near-)essential genes may be enhanced under difficult circumstances, where fitness values across genetic backgrounds converge. In our example, step-wise evolution of Bem1p is still feasible given laboratory conditions (Brauns et al., 2020). Yet, for other proteins, medium quality change may be the only manner to circumvent fitness valleys.

Finally, we determined the key for successful epistasis predictions. As exemplified by the robustness of results to many alternative growth detail formulations, the key here is combining the mesotype with dosage noise. In the case of a sharp mesotype threshold as for polarity, acting on a noisy protein like Cdc42p (about four times the typical coefficient of variation in (Chong et al., 2015)), epistasis with a strong perturbation as with Δ*bem1* makes other mechanistic details irrelevant. Indeed, when examining high-throughput literature data, functional information proved necessary and sufficient to generate meaningful predictions for epistasis (Figure 4D), in line with the ontotype strategy (Yu et al., 2016). As the sharp mesotype implies essentiality (around 19% of yeast genes (Giaever et al., 2002)) or toxicity, generally other mesotypes are more appropriate. Yet, given the simple functions with which fitness landscapes as function of single genes can be fitted (Keren et al., 2016), the mesotype approach still seems feasible to model many biological networks.

Using our findings on yeast polarity as a template, we envision a road-map to apply to general genotype-phenotype maps (Figure 5). The core functional component, in this case polarity, is to be modelled by justifiable coarse-graining, which results in the mesotype of the system. This mesotype in turn emerges from functional subunits (Brauns et al., 2020), identifiable from the rigorous analysis of the underlying biophysics. Currently, yeast polarity stands alone at the ideal intersection between complexity and required mechanistic knowledge. Mesotype generation for other systems would again require detailed numerical analysis of reaction-diffusion systems, or results more simply from dose-reponse curves when spatial information is not essential for the chemical reactions. But once multiple model systems (such as the PAR protein system in *Caenorhabditis elegans* (Goldstein and Macara, 2007)) have been described in this manner, it may be possible to construct a limited library of recurring subunits. Based on this library, construction of the corresponding mesotype may be much simpler. In combination with a coarse-grained view of cell growth and noisy protein production, this paves the general path to bottom-up (population) phenotype prediction.

**Figure 5.**
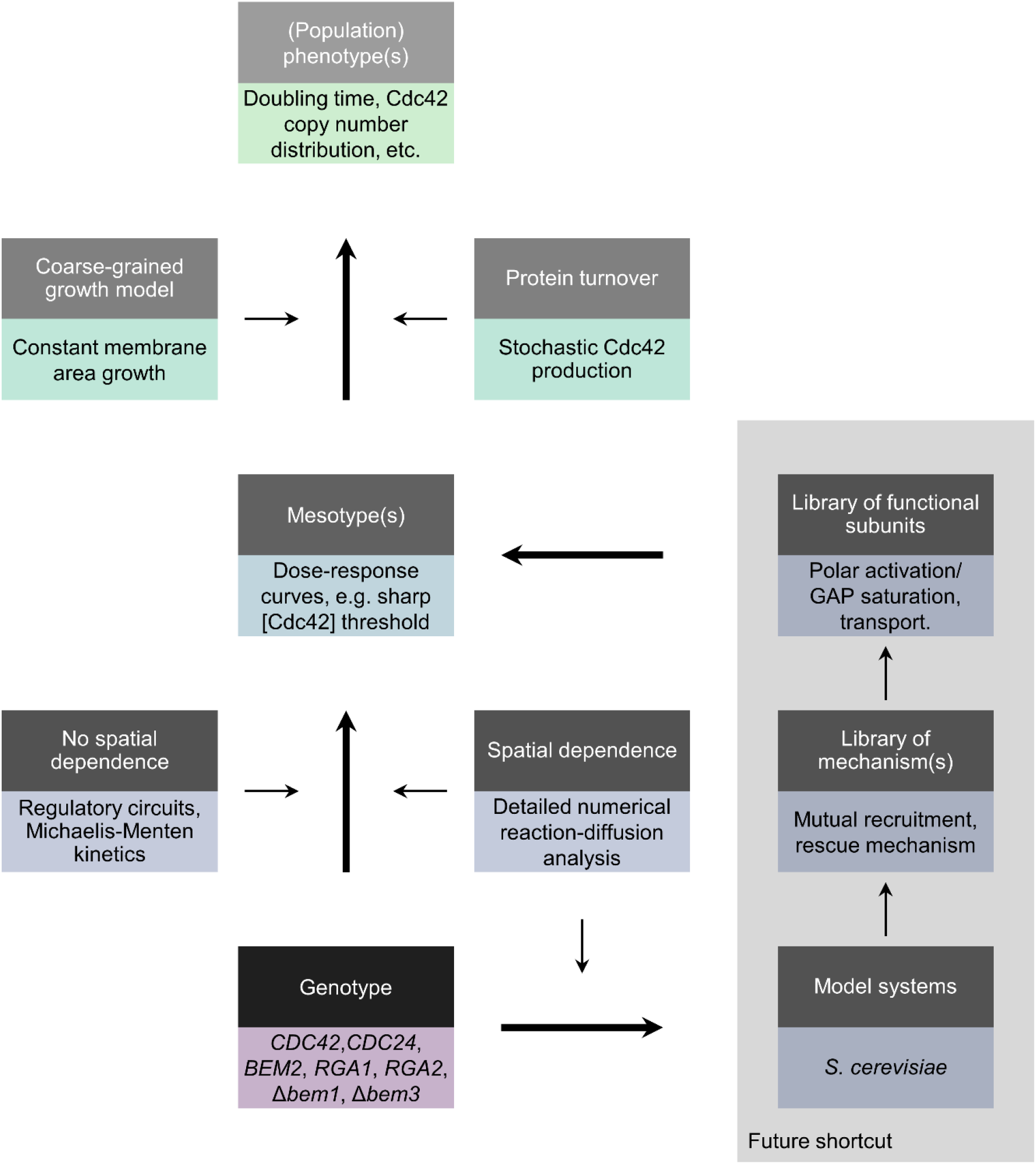
Proposed flow chart for phenotype predictions through intermediate levels. Bottom-up approach for phenotype prediction from genotype through mesotypes, which result from selecting the appropriate functional subunits. The mesotype will typically result from dose-response curves, which are relatively simple to derive from Michaelis-Menten kinetics when there is no spatial dependence. Otherwise, either rigorous numerical analysis of reaction-diffusion equations is required, which may in the future be bypassed when more model systems are analyzed (e.g., polarization in *S. cerevisiae*, the min-system in *E. coli*, PAR-system in *C. elegans*). From a sufficiently large library of functional subunits, the applicable mesotype for the system of interest may be extrapolated. Bottom-half in each box exemplifies the flow chart with the yeast polarity case.

### Limitations of the study

Yeast polarity, our system of interest, is an example of a self-organization network, where we cannot by default assume tractable Michaelis-Menten kinetics, because we cannot neglect the spatiotemporal interplay of protein species. In other model systems where proteins can be assumed to be well-mixed, other modelling approaches have been shown to be highly effective. For example, in budding yeast there is the iMeGroCy model (Pelillo et al., 2018), more than 100 protein species were covered in a mechanistic model of macrophage polarization (Zhao et al., 2021), and modelling metabolism in *S. cerevisiae* can even encompass thousands of protein species (Sánchez et al., 2017).

Although mechanistic information is generally not widely available, this is not always a problem. As we demonstrated in Figure 3A and more specifically in Figure 4D, we extended predictions beyond polarity, where much is known, with mutants that affect the cell in other modules, where less details are known, as for example the case with Nrp1p. This extension is still useful for epistatic predictions if there is at least functional information about the added mutant gene, but becomes dubious once the mutant information is more generic.

Another consideration is to what extent our approach will work for other model systems where a core module depends on the intricate interplay of spatiotemporal interactions. The mesotype relies on coarse-graining abundant, detailed mechanistic information concerning the core biochemical network, and this property currently prohibits application to many other self-organizing systems. Concerns are that some protein networks, particularly for complex organisms, become too complex to coarse-grain, or that including the diverse environmental dependencies required to fully reveal biological functions (Bergelson et al., 2021), also sharply increases complexity. After all, other systems that share the necessary biochemical network knowledge, for example the Min system in *E. coli* (Ramm et al., 2019), exhibit less complexity, while for systems with higher complexity insufficient mechanistic information is available.

Presumably, advances in the near future in systems with insufficient information will follow. For example,, recent theoretical advances provide a promising tractable framework for the PAR-system of *C. elegans* (Geßele et al., 2020). But for more complex networks, our method may take several decades to fully roll-out. For example, the time span between discovery of the first Cdc42p GAP and revealing its mechanism in yeast polarity spans three decades (Brauns et al., 2020; Zheng et al., 1994). The key question on whether the definition of a tractable mesotype following rigorous analysis of the biochemical network remains possible for more difficult networks, is therefore still awaiting a definitive answer.

### Author Contributions

Conceptualization, WD and LL; Methodology, WD; Software, WD; Validation, WD and ES; Formal Analysis; WD and ES; Investigation, WD, ES and LL; Data Curation WD and ES; Writing - Original Draft, WD; Writing - Review & editing, WD, ES and LL; Visualization, WD and ES; Supervision, LL; Project Administration, LL.

## Acknowledgements

We thank Marit Smeets for flow cytometry measurements of two strains. We gratefully acknowledge Leila Iñigo de la Cruz for growth data on RWS1421 and constructing strains yLIC132 and yLIC133. We express our gratitude to Enzo Kingma and Wessel Teunisse for constructing strain yWT3. Additionally, we thank Fridtjof Brauns and Jos Zwanikken for careful reading of the manuscript and Sophie Tschirpke for help in optimizing figure formatting. We acknowledge (Smith et al., 2015) for making the viridis colormap generously available. LL and WD gratefully acknowledge support from the Netherlands Organization for Scientific Research (NWO/OCW), as part of the Gravitation Program: Frontiers of Nanoscience. LL and ES gratefully acknowledge funding from the European Research Council (ERC) under the European Union’s Horizon 2020 research and innovation programme (Grant agreement No. [758132]).

## Materials and Methods

## RESOURCE AVAILABILITY

### Lead contact

Further information and requests for resources and reagents should be directed to and will be fulfilled by the Lead Contact, Liedewij Laan.

### Materials availability

Strains generated in this study are available from the Lead Contact on request, we will require a completed Materials Transfer Agreement due to TU Delft policy.

### Data and code availability

The datasets and code generated during this study are available as Supplemental Information.

## EXPERIMENTAL MODEL AND SUBJECT DETAILS

### Microbe strains

All strains used in this study are *S. cerevisiae* strains in the W303 background. A full list of strains used in this study can be found in the Materials and strains table. Cells are kept for long-term storage in a frozen glycerol stock at -80°C, and are cultured for experiments at 30°C by inoculating in liquid fresh media. Strains RWS116 and RWS1421 require absence of methionine in the media for growth, so we used YNB (0.69% w/v, Sigma-Aldrich) with CSM -MET (0.77% w/v, Formedium) amino acid mix and 2% dextrose (Sigma-Aldrich). For fluorescence microscopy of yLL129a (Figure 1B), we used low-fluorescence media, namely non-fluorescent nitrogen base (0.69% w/v, Formedium), CSM amino acid mix (0.79% w/v, Formedium) and 2% dextrose. For the growth rate assay in Figure 4B, we used YPD (10 g/L Yeast extract, 20 g/L peptone, 0.1/0.5/2% dextrose, 20 µg/ml adenine) and 0.25 mM auxin for strains yLIC133 and yWT3.

## Method details

### Extended explanation cell cycle model

#### Size expansion

From (Goranov et al., 2009), two growth phases are distinguished, with seemingly linear growth of volume with respect to time for the mother and sublinear volume growth for the bud, possibly consistent with constant growth of membrane area across time. Two phases are similarly visible in (Ferrezuelo et al., 2012), but also indicating sublinear dependency between volume and time. In light of absence of consensus, we accommodated both the constant area and constant volume expansion across time implementation in our model, but used the former by default. However, for the description of GAP epistasis this choice is inconsequential, see Figure S2. We follow the observation in (Ferrezuelo et al., 2012) that mother expansion stops during bud growth. We hence describe the size expansion as:

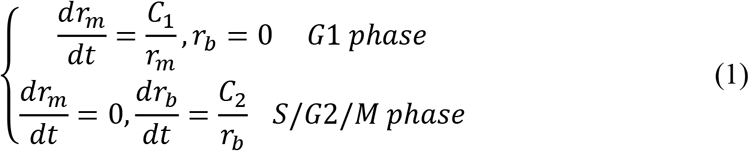

where *r_m_* and *r_b_* are mother and bud radius respectively, and *C_1_* and *C_2_* are constants.

To set *C_1_*, we assume an optimized WT such that at checkpoint 1, the minimum size requirement *r_m_ ≥ r_min_* is typically met at the same time that the minimum G1 time requirement (time before polarization starts must be *≥ t_G1,min_*) is met, and that subsequent polarization time is minimal. After G1, the (squared) mother radius is then (integrating equation 1 from 0 to *t_pol,min_*, the duration needed to polarize for a fast WT):

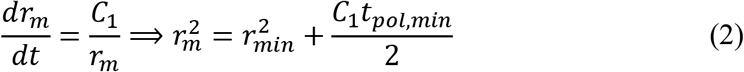

From (Ferrezuelo et al., 2012; Goranov et al., 2009), we infer final bud volumes to be around 70% of mother volumes. This means the bud radius after the next M-phase *r_b_*=0.7^1/3^ *r_m_* ≈0.89 *r_m_*. For self-consistency, this new cell must then expand to size *r_min_* again at checkpoint 1, such that:

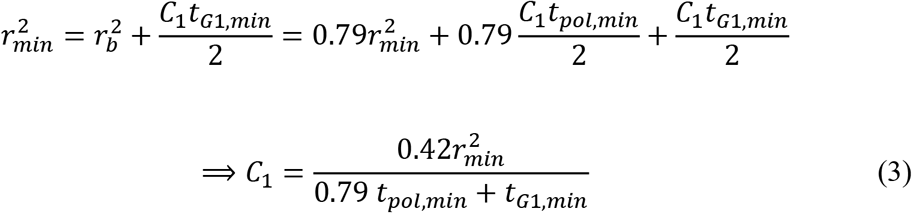

From Table S6 of (Ferrezuelo et al., 2012), we see the volume at Start in glucose of WT is 36.1 µm^3^, which assuming a spherical cell implies *r_min_* ≤ 2.05 µm, which we round to 2 µm. From Table S7 of the same paper, we see Time at Start for mother WT cells to be 15.6 min., so we set *t_G1,min_* to this value. Note it is this value that alters when we incorporate the *nrp1* deletion. Finally, we estimate *t_pol,min_* from (Howell et al., 2012), where it is noted that polarization of Bem1p typically occurs in 2 minutes starting from the point of first detection of significant fluorescence. To estimate the full minimal polarization time including the time when Bem1p accumulation is still too faint, we turned to the power spectrum of the oscillation time in Figure 1F of that paper, which peaks around 0.2 1/min., corresponding to a time to build a spot of 5 min., so we set *t_pol,min_* to this value.

From these parameter values, we calculate *C_1_*=0.086 µm^2^/min from equation 3. This value is consistent with the aforementioned papers of (Ferrezuelo et al., 2012; Goranov et al., 2009). For example, reading out from the plot in Fig. 7C of (Goranov et al., 2009) we see for the mother growth phase a shift from 1500 voxels to 1700 voxels in -30 to 0 min. 1 voxel is 0.26 *μm*^3^, so

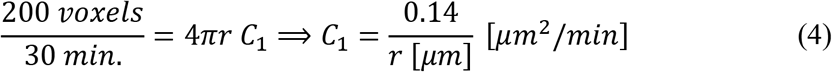

With a typical size of 2 *μm*, we have *C_1_* ≈ 0.07 μm^2^/min. For (Ferrezuelo et al., 2012), we see volume growth of 0.15 μm^3^/min, and given 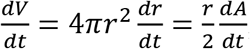 we expect *C_1_* ≈ 0.15 μm^2^/min. Therefore, our value of *C_1_*=0.086 μm^2^/min is within the expected range.

The value of *C_2_* (equation 1) follows from calibration, by running the model implementation (see further on this Materials and Methods) and varying *C_2_* such that under WT parameters, the doubling time is close to 83 minutes, as expected from literature (Laan et al., 2015). This calibration yields *C_2_=*2.1324 µm^2^/min. Again, this is in line with literature, as the rate during bud growth phase is roughly twice as fast as the isotropic growth phase in (Ferrezuelo et al., 2012; Goranov et al., 2009). To accommodate different growth media into the calculation as for the growth rates of Figure 4A, both membrane growth rate can be multiplied by the same factor. For Figure 4A, this factor was between 0.3 and 1 (steps of 0.1).

We also set a maximum size value, *r_max_*, which plays a role when Cdc42 turnover has been unfavorable to prevent polarization. The value is set at 6 µm, which prevents excessive calculations on cells with very poor forecasts, and accommodates the smallest and thus healthiest >90% of the cells of the ill Δ*bem1* background, assuming a normal distribution for the cell diameter with mean 6 µm and standard deviation 4.3 µm, with values estimated from Figures 1E and 1F of (Laan et al., 2015). Setting the maximum radius to i.e. 20 µm has a negligible influence (<0.25%) on simulated doubling times of WT and Δ*bem1* cells.

### Cdc42p turnover

We expect Cdc42p expression bursts at intervals and with sizes which are exponentially distributed (Friedman et al., 2006). In this paper, it was also explained how the average burst interval time and size can be inferred from flow cytometry data. The former, *t_b,WT_*, follows from dividing the average cycle time by the shape parameter of a gamma fit on the endogenous Cdc42p distribution across the population. The latter, *p_b,WT_*, in principle follows from dividing the average Cdc42p copy number which can be estimated from (Kulak et al., 2014) to be 8700, by the same shape parameter of the aforementioned gamma fit. In this way, the mean of the gamma distribution should equal 8700. However, as we explicitly wish to include degradation into our model to better mimic the critical part of the turnover concerning passage of the mesotype threshold, we must adjust the scaling parameter slightly. Following calibration as in the previous section, we adjusted *p_b,WT_* (see Table S1) such that the average copy number of Cdc42p in WT is close to the 8700 from (Kulak et al., 2014). This required an 18,67% reduction of the original *p_b,WT_*.

The flow cytometry for setting *t_b,WT_* and *p_b,WT_* was performed using strains RWS116 and RWS1421, from (Freisinger et al., 2013; Gulli et al., 2000)). RWS1421 contains C*DC42pr-GFP-CDC42* in addition to its endogenous *CDC42*, as this GFP tagged Cdc42p is not fully functional. However, we do not expect this tag to be of major influence on the inferred copy number from fluorescence, as the main issue is mislocalization of this Cdc42-variant (Watson et al., 2014). RWS116 has the same strain background as RWS1421 but without fluorescence, to serve as a background measurement. Fluorescence measurements were acquired using FlowJo CE software and performed on a BD FACScan flow cytometer. Cells were pregrown in YNB (Sigma) + CSM -Met (Formedium) + 2% dextrose (Sigma-Aldrich), diluted to an OD*_600_* of 0.1 and measured after 15h.

The flow cytometry data was then analyzed (for the script see SI experimental code) by manually imposing a polygon in the Forward/Side scattering plot to remove data points that likely correspond to debris or dead cells (gating). Using Matlab’s R2016a built-in *mle* function, we fit a gamma distribution using maximum likelihood, to the background and endogenous Cdc42p distribution. This yields the shape parameters *k* = 2.36 ± 0.05 and 4.74 ± 0.07 and the scale parameters *θ* = 20.2 ± 0.5 and 59.6 ± 0.9 for the background and the endogenous distribution respectively, where the ± sign reports the 95% confidence interval.

We then approximate the analytical deconvolution of the endogenous distribution by setting the deconvolved distribution as a gamma distribution (Stewart et al., 2007), where parameters follow from subtracting the mean and variance of the background from the mean and variance respectively of the endogenous signal. In that paper, the gamma approximation for deconvolution is shown to hold as long as the scaling parameters of the background and endogenous signal are within an order of magnitude, as is the case here. We retrieve for the background corrected endogenous signal the shape parameter *k* = 3.48 ± 0.07 and the scale parameter *θ* = 67.6 ± 0.9. Here the 95% confidence interval indicated by the ± sign follows from 2.5% and 97.5% quantiles of Monte Carlo simulations (10000 runs) of calculations of the shape and scaling parameters assuming normal errors on the background and uncorrected endogenous parameters, also assuming independence between a draw of a shape parameter and a draw of a scaling parameter.

The growth rate assay for RWS1421 was performed in the same media type as the flow cytometry, in an Infinite M-200 Pro Tecan plate reader at 30°C.OD*_600_* measurements (9 nm bandwidth) were conducted with an interval of 12 minutes for a total duration of 24 hours, using a Thermo Fisher Scientific Nunclon 96 Flat Bottom Plate input template. After an initial 1000s of orbital shaking (1 mm amplitude), linear shaking between measurements (25 flashes, 5 ms settle time) lasted 330s each time (amplitude 1mm).

Only three wells (E2, E3, E4) were designated for RWS1421 in the appropriate medium. These were analyzed using a home-written Matlab GUI already used in (Brauns et al., 2020) and made available as Supplemental Information. The GUI allows the user to import the plate reader output file in excel format, after which the user can specify a time bandwidth within which to perform an ordinary (OLS) or weighted least squares (WLS) fit (OLS/WLS regressions described in (Heij et al., 2004)). This fit follows from a linear regression on the log (base 2) of the (OD*_600_* – background OD*_600_* value), where the background is set as the mean of the first ten OD*_600_* values in that well. The weights for WLS are set by the reciprocal of the difference between log of the OD*_600_* values with the instrument error added and subtracted. The instrument error is approximated by taking the standard deviation of the first ten OD*_600_* or by 10^-digit^/2 where digit is the value of the exponent with base 10 of the last significant digit. The latter case applies when the data output from the plate reader has rounded the data to an extent that the first ten point are identical. WLS is performed for every fit window of a user-defined size that fits within the time bandwidth that the user also defines, and that only contains data corresponding to the longest time span above a user defined signal-to-noise ratio (with noise from the instrument error). For every fit window, the reciprocal of the doubling time is the slope of the WLS regression, which follows a Student’s t-distribution with the number of data points in the fitting window minus 2 as the degrees of freedom (Heij et al., 2004). This distribution sets the standard error. The fastest doubling time which pertains to a fit with a R^2^ squared above a user defined value, is chosen to be the doubling time.

Since the flow cytometry measurement protocol suggests the protein distribution reflects more accurately as the distribution is during late log phase (measuring 15h after dilution), we intentionally fit above an OD above 0.1, roughly between 10 and 14h into the growth experiment. We set our fitting window size to 21 points, weighted least squares option to on, minimum R^2^ to 0.9 and mininmal signal-to-noise ratio to 2. The doubling times following the analysis in the GUI were 129, 199 and 199 minutes. We take the median value and round to 200 minutes to use as the cycle time for the calculation of *t_b,WT_*.

Table S4 contains the flow cytometry (raw, processed and fitted) of RWS116 and RWS1421 used in this study, see also Figure S3. Table S5 contains OD*_600_* growth assay measurements of RWS1421 (with fits).

### Polarity mesotype

The rigorous analysis of the reaction-diffusion equations and numerical simulations done in (Brauns et al., 2020) provided the support to simplify the a priori complex dependency of polarization success on polarity protein concentrations and reaction rates to a simple cone-like phase space (see Figure 1D). More concretely, this also resulted in the support to justify a simple approximation for the polarization time, see the qualitative plots in Figures 23 and 24 in (Daalman, 2020) (courtesy of Fridtjof Brauns). We approximate the polarization time *t_pol_* as an exponential:

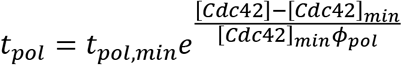

where [Cdc42p]*_min_* is the polarity mesotype and *φ_pol_* is a scaling parameter to govern the theoretically predicted exponential increase in polarization time when there is excessive Cdc42p. As the aforementioned qualitative plots indicate similar behavior for low GAPs concentrations (relative to Cdc42p), we assume [Cdc42p]*_min_ φ_pol_* for both the *BEM1* and Δ*bem1* background. As no clear deleterious effects are apparent for high induction of Cdc42p under the Gal1 promoter in (Brauns et al., 2020), we set

*ϕ*_*pol*_ at a sufficiently high value of 25 for the Δ*bem1* background. For the *BEM1* background, we notice that setting a value of 500 yields the approximately self-consistent result [Cdc42p]_min_,Δb*_em1_ φ_pol_* Δb*_em1_* = 116 ·25 ≈ [Cdc42p]*_min,BEM1_ φ_pol BEM1_* = 4 ·500.

Because of the shape of the phase space, the mesotypes are simple linear functions of GAP concentrations, such that we only require to fit the polarity mesotype for the WT and Δ*bem1* background and fit relative shift in mesotype for a *bem3* deletion and for the *bem2* deletion (the same in both backgrounds). The relative shift in mesotype with both GAPs deleted is then simply the sum of the two relative shifts, and is not modelled to alter upon *nrp1* deletion. Thus, four mesotypes (and a change in *t_G1,min_* for the *nrp1* deletion) describe all polarity mutants from (Laan et al., 2015) considered in this study.

As a final note, we impose for practical purposes a maximum polarization time of 600 min., to truncate excessive long polarization times. Extending this time to 2000 min. has no influence on doubling times of WT and Δ*bem1* cells.

### G1 timing

We also experimented with a more realistic (and less coarse-grained) modification of the modelled cell cycle progression to improve the quantitative match of the *BEM1-NRP1* epistasis in Figure 3A. Suppose for example the *Δbem1* cells if the assumed minimum G1 time set is not a constant but results from a distribution. This is more realistic, as times for symmetry breaking in daughter cells can exhibit notable stochasticity (Moran et al., 2019). Some Δ*bem1* cells have an early opportunity to fulfil the mesotype threshold concentration requirement, with which they usually struggle, while others are delayed more. This increases the cell-to-cell variation in fates in G1, since cells with fast G1 times are most likely to generate a first spot, while slow cells never generate this spot and thus do not show up in the statistics. This is how less coarse-graining can lead to a larger decrease in G1 times from WT to Δ*bem1* than is the case with constant G1 waiting time *t_G1,min_*. Implementing a minimum G1 time distribution yields a slight quantitative improvement for the G1 times in the genotype trajectory of Figure 3D (see Table S2). Thus, other unmodelled parts of the cell cycle are likely responsible for the remaining quantitative discrepancy in describing G1 times.

### Fitting parameters cell cycle model

Doubling times of (Laan et al., 2015) in Figure 3A were fitted using the built-in *fminsearch* on a normalized score objective for varying [Cdc42]*_min_* and manual inspection for setting *t_mut_* (to 0.75) for the *nrp1* deletion. The objective involves the sum of squares of the fitting errors divided by the standard errors of the experimental literature values. The experimental error for the Δ*bem2* Δ*nrp1* is not available, and conservatively set at 30 minutes.

### Computational implementation cell cycle model Overview algorithm

Cell cycle model simulations were performed in MATLAB R2016a following a partial leap-like Gillespie algorithm (Gillespie, 2001) implementation (the G1 time until *r<r_min_* and *t_1_<t_G1,min_*, *t_pol_* and the time through S/G2/M are one leap each). Cell cycle model parameters are summarized in Table S1. The core function Mesotype_model.m, function to generate the data for the figures and an example script to demonstrate the functionality are found as Supplemental Information. Information considering the input parameters for each function are included in the code, and can be called from Matlab’s command window by e.g., the command “help Mesotype_model”. A pseudocode representation of the core Mesotype_model.m function follows the descriptions in the next paragraphs.

An initial population (ancestor seed) asynchronized across a bandwidth of 83 minutes (all cells with equal radii of 2.2 µm and without proteins) is defined. For every iteration in the simulated growth loop, the lowest 50% of the population in terms of simulation time since the ancestor seed is considered for growth, to avoid excessive asynchrony for the cells. Each of those cells aim to progress through the cell cycle, where the step to fulfill all polarity conditions (*r<r_min_* and *t_1_<t_G1,min_* and [Cdc42p]>[Cdc42p]*_min_*) is the only step where permanent cell death can take place. The 50% of the cells not considered in the iteration are temporary scored as death, and this is reset in every iteration.

The population grows until a user-defined population size placed as an input argument in the core function, after which the colony is diluted to 1000 cells. Throughout this study, we use a population size value of 5 million except for the single cell time trace where this was set at 2e3 to avoid diluting too many of the initial cells. After dilution, the population grows again until the same user-defined population size, or the highest user-defined time for copy number and volume distribution probes if this requires the simulation to continue. All cells left alive must have a simulated time since the ancestor seed to consider a copy number or volume probe complete.

If the colony fails to complete the simulation due to all cells dying prematurely, the simulation is restarted. If the colony fails five times, the simulation returns a default doubling time of 1e9 minutes. Otherwise, doubling times are the average of the last hundred moving window (size 201 min.) linear regressions on the log number of cells.

### Pseudocode representation of our physical cell cycle model

**% Initialization**

num_cells = 1000, radius=2.2, copy_number =0, simulation_time = -83 to 0

num_tries = 0

while num_cells < 5e6 & num_tries < 5

for all alive cells

**% Isotropic growth until *r* = *r_min_* and at least time_step > *t_G1,min_***
time_step = max( 4π(*r_min2_* - *r^2^)* / *C1, t_G1,min_* )
num_bursts ∼ poiss( time_step / *t_b,WT_* ) % draw form poisson distribution distribute burst_times uniformly across time step
for all bursts

burst_size ∼ exp(*p_b,WT_*) % draw form exponential distribution
burst_size = burst_size * exp(burst_time / *τ_h_*) % correction for dilution
mother copy number = mother copy number * exp(time_step / *τ_h_*)
% correction for dilution
mother copy number = mother copy number + sum of all burst sizes
mother radius = mother radius + *C_1_* time_step % update mother radius
**% Isotropic growth until mother copy number > *P_min_* or *r* > *r_max_***
**while mother copy number < *P_min_* and *r* < *r_max_***

time_step ∼ exp(*t_b,WT_*) % draw form exponential distribution
burst_size = ∼ exp(*p_b,WT_*) % draw form exponential distribution
mother copy number = mother copy number * exp(time_step / *τ_h_*)
% correction for dilution
mother copy number = mother copy number + burst_size
mother radius = mother radius + *C_1_* time_step % update mother radius
end
for all cells with **mother copy number** < *P_min_* and *r* > *r_max_*

mark cells as dead
for all alive cells

**% Isotropic growth until polarization is achieved**
time_step = *t_pol,min_* exp( (mother copy number - *P_min_*) )/( *P_min_ ϕ_pol_* ) )
num_bursts ∼ poiss( time_step / *t_b,WT_* ) % draw form poisson distribution
distribute burst_times uniformly across time step
for all bursts

burst_size ∼ exp(*p_b,WT_*) % draw form exponential distribution
burst_size = burst_size * exp(burst_time / *τ_h_*) % correction for dilution
mother copy number = mother copy number * exp(time_step / *τ_h_*) % correction for dilution
mother copy number = mother copy number + sum of all burst sizes mother radius = mother radius + *C_1_* time_step % update mother radius update mother simulation_time
**% Polarized growth of bud**
time_step = 4π ( (0.7)^1/3^ mother radius)^2^ / C*_2_*
num_bursts ∼ poiss( time_step / *t_b,WT_* ) % draw form poisson distribution
distribute burst_times uniformly across time step
for all bursts

burst_size ∼ exp(*p_b,WT_*) % draw form exponential distribution
burst_size = burst_size * exp(burst_time / *τ_h_*) % correction for dilution
mother copy number = mother copy number * exp(time_step / *τ_h_*)
% correction for dilution
mother copy number = mother copy number + sum of all burst sizes
bud radius = bud radius + *C_2_ \** time_step % update bud radius
bud copy_number = copy_number * bud volume / total volume % update bud
copy_number
Define buds as new cells
if no cells left

num_tries = num_tries + 1
% Re-initialize
num_cells = 1000, radius=2.2, copy_number =0, simulation_time = -83 to 0
continue with next iteration while-loop
if num_cells > 5e6 & dilution did not occur yet

Dilute: Randomly select 1000 cells and restart while loop
end

Note: checks for simulated measurements of volumes and copy numbers occur in each time_step. Required intermediate radii follow deterministically and are stored. Intermediate copy numbers follow from uniformly distributing the bursts across the time_step and store the intermediate copy number update of the bursts that fall in the time until the simulated measurment.

### Growth assay varying medium quality

Strains for the growth assay of Figure 4B were inoculated in YP2%D + 20 µg/ml adenine from glycerol stock in glass tubes. These were then grown in a turning wheel for at least 24h at 30 °C. After this pre-growth the cultures were diluted (at least 100x) to an OD*_600_* of approx. 0.05 into a 96-well plate. Each well contains 100 µL of one of the three medium types with differing glucose concentrations (YP + 0.1%, 0.5%, and 2% dextrose). Auxin was added to a final concentration of 0.25mM this medium to induce the Bem1p degradation. 0.25mM auxin was also added to the wells with WT yeast as a control. Then the OD*_600_* was measured for 48h using a Biotek Epoch™ 2 Microplate Spectrophotometer using linear and orbital shaking at 30C, as described in (Brauns et al., 2020)

### Predictions using literature data

Interaction and phenotype data for Figure 4D were obtained from BioGRID (Stark, 2006) and SGD (Cherry et al., 2012) respectively (date of access March 2018), see also Table S6 for detailed references per interaction. Concretely, interactions with Bem1 are from https://thebiogrid.org/32897/summary/saccharomyces-cerevisiae/bem1.html. Data for competitive fitness mutants are from the SGD Project, https://www.yeastgenome.org/observable/APO:0000110 [6 March 2018]. Data for fermentative growth mutants are from the SGD Project. https://www.yeastgenome.org/observable/APO:0000308 [6 March 2018]. Data for critical cell size at START (G1 cell-size checkpoint) mutants are from the SGD Project. https://www.yeastgenome.org/observable/APO:0000141 [8 March 2018]. Data for cell cycle progression in G1 phase mutants are from the SGD Project. https://www.yeastgenome.org/observable/APO:0000255 [8 March 2018]. Data for genes annotated for "proteasome" are from the SGD Project. http://yeastgenome.org/search?category=locus&geneMode=wrap&page=0&q=proteasome [8 March 2018]. Data for genes annotated for "ribosomal" are from the SGD Project. http://yeastgenome.org/search?category=locus&geneMode=wrap&page=0&q=ribosomal [8 March 2018]. Data for Genes annotated for "phospholipid" are from the SGD Project. http://yeastgenome.org/search?category=locus&geneMode=wrap&page=0&q=phospholipid [8 March 2018].

We consider all ’Positive Genetic’ and ’Synthetic Rescue’ annotations as positive interactions, and ’Negative Genetic’ and ’Synthetic Lethality’ as negative interactions. For the competitive fitness mutants, we only considered those with phenotype competitive fitness: decreased (negative interactions) or increased (positive interactions) in YPD. For the fermentative fitness mutants, we only considered those with phenotype competitive fitness: decreased rate (negative interactions) or increased rate (positive interactions) in YPD. For the mutants with altered critical cell size at START (G1 cell-size checkpoint), we only considered the null, reduced and overexpression mutants. For the overexpression mutants, we assumed the null/reduced mutants have the opposite effect (e.g., decreased size to increased size). This was done similarly for the cell cycle progression in G1 phase mutants, where we considered the same mutant types but for the phenotypes cell cycle progression in G1 phase: delayed/increased duration/increased rate/decreased duration. When interactions in all previously mentioned categories led to a conflict between effects (sources have opposing opinions), we either choose the effect that is most commonly reported or discard the gene if there is no single most commonly reported effect. We overwrite the genetic interaction with Bem2 as positive (Laan et al., 2015).

We assume for simplicity that all genes annotated for “proteasome” have a negative effect on degradation when mutated. We assume for simplicity that all genes annotated for “ribosomal” have a negative effect on translation and hence protein production when mutated. We assume for simplicity that all genes annotated for “phospholipid” have a negative effect on phospholipid synthesis and hence membrane area growth when mutated. Similar to the fitness mutants in the coarse phenotype mutant set, membrane area growth rate mutations are incorporated in *C_2_* alone, as this modification is more effective in changing fitness for the WT background than changing both *C_1_* and *C_2_*.

### Strain construction

Firstly, the *ade2* deletion was performed through transformation by homologous recombination. The repair fragment was generated by overlap extension PCR using primers olic24, olic15, olic18 and olic26, and encompasses the *URA3* (from a template derived from the pRL368 plasmid (Wedlich-Soldner, 2004)) gene with homology regions with *ADE2* (from a yLL3a genomic template). Selection on uracil dropout media completed the transformation of yLL3a to yLIC132. Similarly, overlap extension PCR using primers olic24, olic20, olic26 and olic21 yielded the repair fragment to remove the URA3 marker by transformation, yielding yLIC133. This strain then formed the basis for yWT3 used in this study, which was generated as follows.

Firstly, the PCR product to integrate *osTIR1* in the HO locus was amplified from plasmid pOsTir1w/oGFP. Plasmid pOsTIR1w/oGFP was a gift from Matthias Heinemann (Addgene plasmid # 102883 ; http://n2t.net/addgene:102883 ; RRID:Addgene_102883. Secondly, using Gibson assembly. a *BEM1* homology region was added to a pG23A plasmid backbone, upstream and in frame of the *mCherry-AID* sequence. Plasmid pG23A was a gift from Matthias Heinemann (Addgene plasmid # 102884 ; http://n2t.net/addgene:102884 ; RRID:Addgene_102884). Further downstream, the *HPHMX6* cassette and another *BEM1* homology region was added, to allow generation of a PCR fragment to integrate the *mCherry-AID* C-terminally on *BEM1* by selecting with Hygromycin at transformation (for primers see Table S8). Starting from strain yLIC133, successive transformation with the PCR fragment from the *osTIR1-KANMX4* plasmid (selecting with G418) and the *BEM1-mCherryAID-HPHMX6* plasmid (selection with hygromycin) generates strain yWT3 (deposition of latter plasmid at Addgene in progress).

## Quantification and statistical analysis

### Pooling and testing growth data

Three experiments with two plates each were done for the glucose sensitivity assay. In each plate we used one and two biological replicates for the *BEM1* and *BEM1-AID* background respectively, with at least two technical replicates per medium. The doubling time of each well was calculated using the Matlab GUI as explained above (section Cdc42p turnover). Per run, we averaged the fitness (defined by the reciprocal of the doubling time) of *BEM1* replicates. We then divided the fitness of each *BEM1-AID* strain by the mean *BEM1* fitness of that run. We then pooled all *BEM1-AID* replicates per medium for data analysis. Wells where no growth were observed were excluded. We neglected the error from the doubling time fits as the largest error source is the variation across replicates. We used a one-sided Welch t-test (Matlab R2016a’s native ttest2, for unequal variances) to test for significant differences, and applied a Holm-Bonferroni correction factor (Holm, 1979) to the significance values for the three t-tests performed.

We disregarded the 0.25% and 1% medium type, to only plot the media containing the most runs. We excluded the other *BEM1* background strain measured in some plates as it lacked the *ade2* mutation present in the *BEM1-AID* strains. Finally, we excluded the biological replicate yWT3a, as its behavior was less comparable to the other two *BEM1-AID* biological replicates. The latter two exclusions do not affect the conclusions on which fitness differences were significant.

### Epistasis predictions with Δ*bem1*

From the interaction sets described in the previous section, we calculated posterior distributions for the probability *θ* of a positive interaction with *BEM1*, using the frequency of such an interaction encountered in literature. We describe the probability of finding a positive interaction given a total number of interactions as following from a binomial likelihood. If we then assume a beta distribution beta(*α*,*β*) as our prior (more specifically, the uninformed prior with beta(1,1)), the resulting posterior distribution for the probability of having a positive interaction *θ* is also a beta distribution, but with parameters *α*+*k* and *β*+*N*, where *k* is the encountered positive interactions and N the number of encountered negative interactions with *BEM1* (Hobbs and Hooten, 2015). The positive and negative interaction counts for this study can be found in Table S7. This gives us the distributions for *θ* for the pooled deleterious fitness mutants (competitive and fermentative), pooled beneficial fitness mutants, mutants that are small at G1 or fast through G1, mutants that are large in G1 or slow through G1, mutants that have less degradation or phospholipid production, and finally mutants that have reduced protein production.

From 10000 random draws of every *θ*, we can simulate the distributions of the differences between pairs of *θ*. In particular, we consider the difference distribution of *θ* between deleterious and beneficial mutants, the difference distribution of the small/fast in G1 mutants and large/slow in G1 mutants, and the difference distribution of the low degradation/lipid and low production mutants. The probability of a difference above zero corresponds to the probability that positive epistasis is more prevalent in deleterious than in beneficial mutants, more prevalent in small/fast in G1 mutants than in large/slow mutants, and more prevalent in low degradation/lipid than in low production mutants. This probability is plotted in Figure 4D. Alternative, we can rewrite this probability as a Bayes factor for the model hypotheses that the positive epistasis is indeed more prevalent in each of the three aforementioned cases. The Bayes factors are these probabilities divided by 1 minus these probabilities, and we turn to (Kass and Raftery, 1995) to interpret these factors.

## Materials and strains table

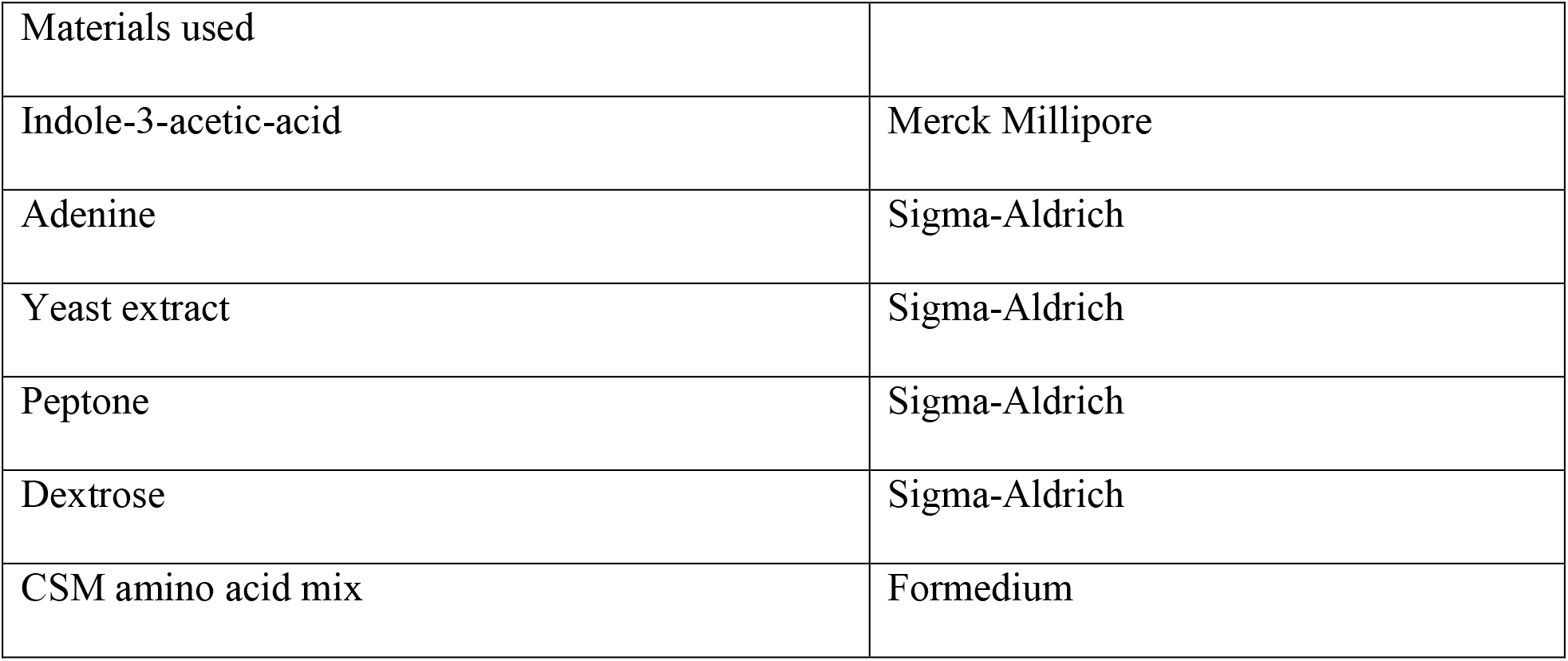
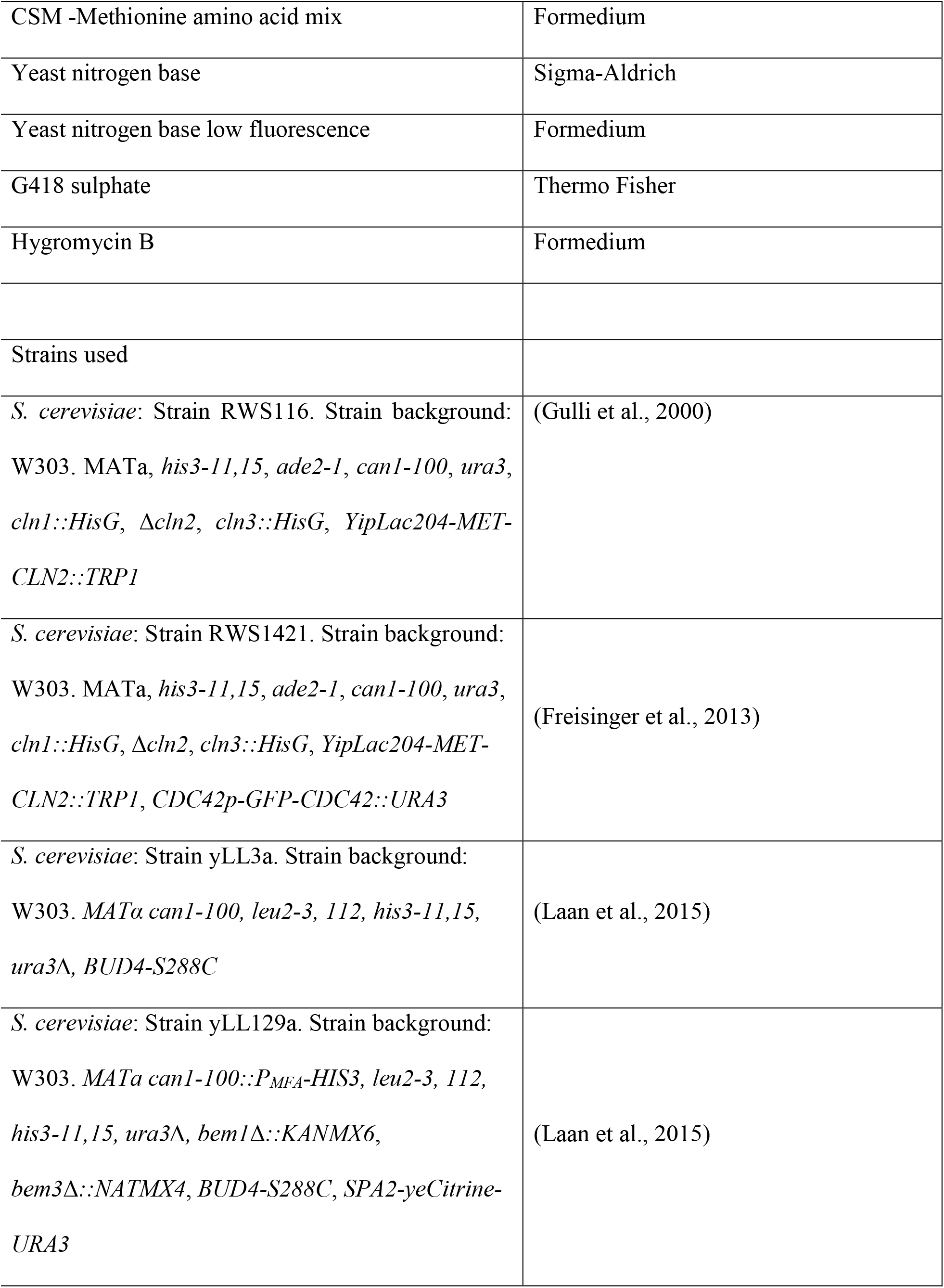
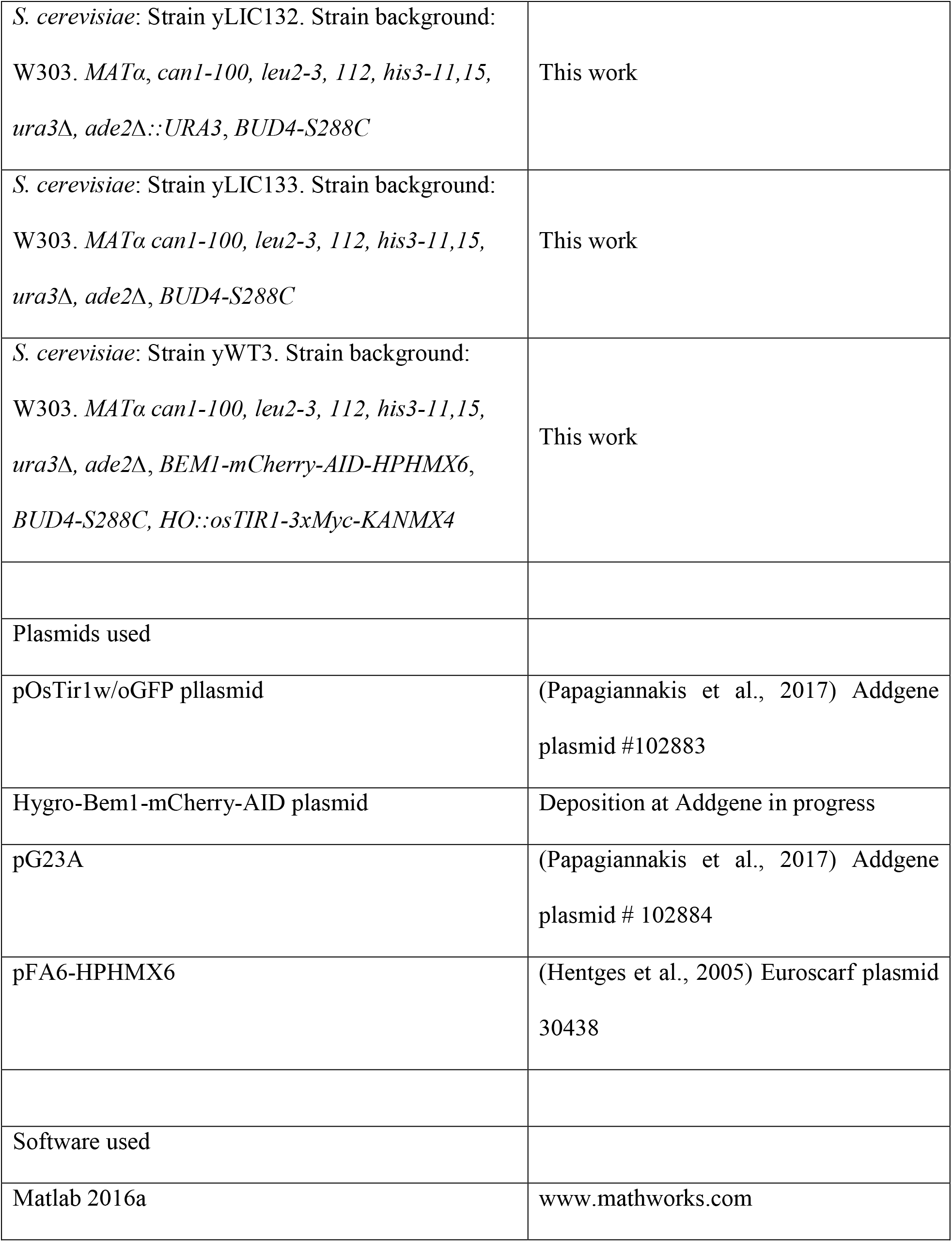
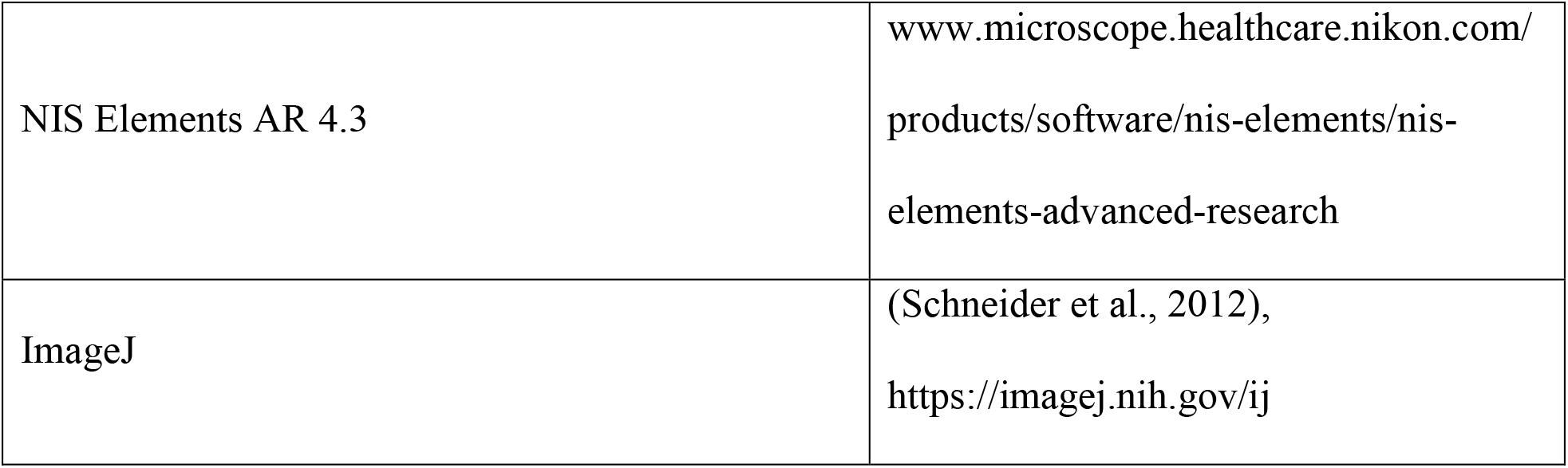

## Supplementary information

**Figure S1.**
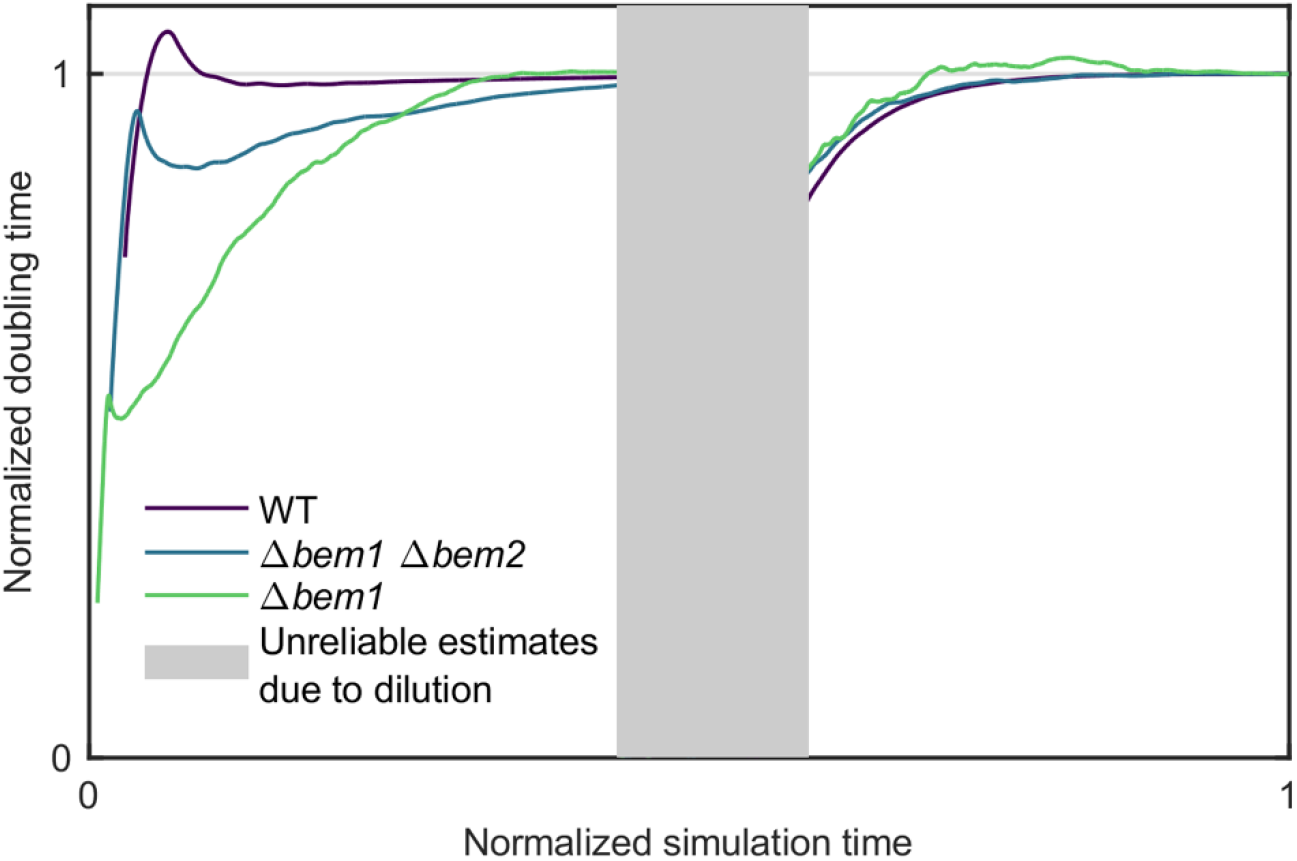
Convergence of simulated doubling times. Doubling times as function of simulation time for a fast (WT), medium (Δ*bem1* Δ*bem2*), and slow (Δ*bem1*) growing strain background. Doubling times and simulation times normalized to their respective final value. The dilution step midway temporarily causes unreliable estimates and these are omitted from display.

**Figure S2.**
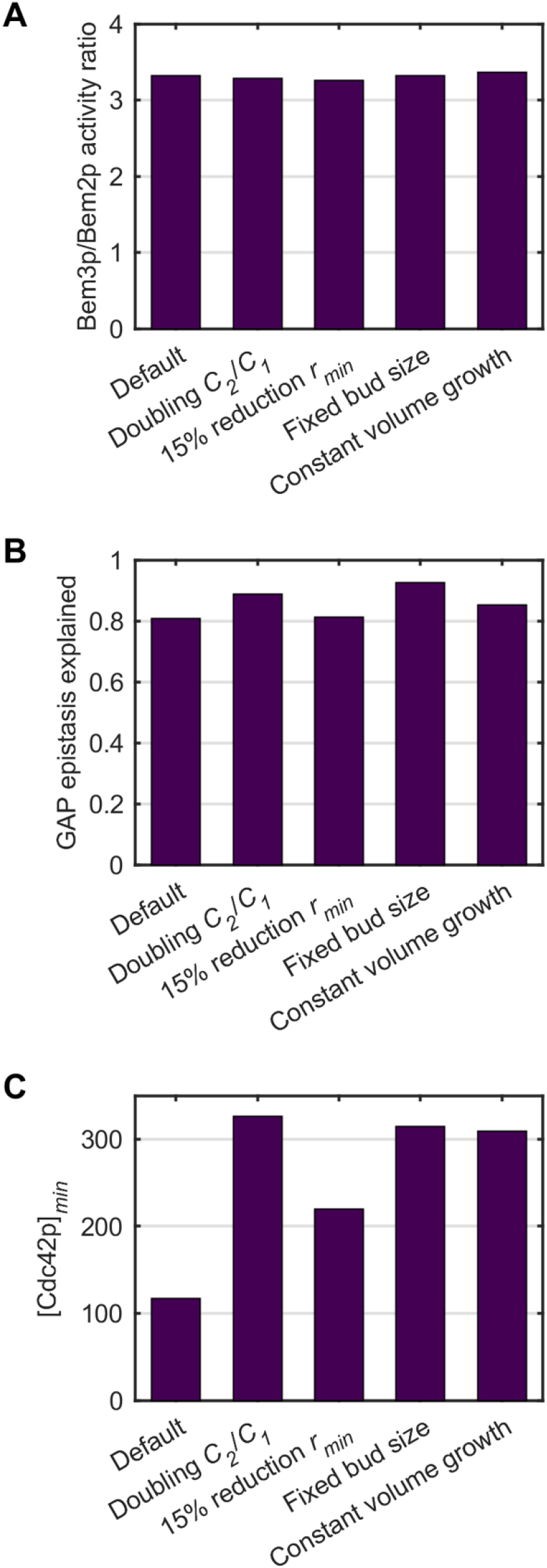
Cell cycle model details effect on relative and absolute quantities. (A) Calculated GAP activity given cell cycle model fits and abundancies from (Kulak et al., 2014), (B) relative multiplicative epistasis (definition of (da Silva et al., 2010)) for the GAPs in the Δ*bem1 NRP1* background and (C) fitted mesotype threshold (minimum Cdc42 concentration to polarize) in the Δ*bem1* background for the default parameter set in the cell cycle model and four variations; doubling of membrane area growth rate *C_2_*, 15% reduction of *r_min_*, change of second checkpoint to a fixed bud size threshold of 1.8 µm, and constant cell volume instead of area expansion. WT membrane growth rates are recalibrated in each case to match 83 minutes. Fits are performed on the doubling times of *NRP1* strains in (Laan et al., 2015) alone, in slight contrast with fits in Figure 3A, to focus on variation due to GAP deletions. Quantities in panels (A) and (B) are relative, dimensionless quantities, whereas the quantity in (C) is absolute.

**Figure S3.**
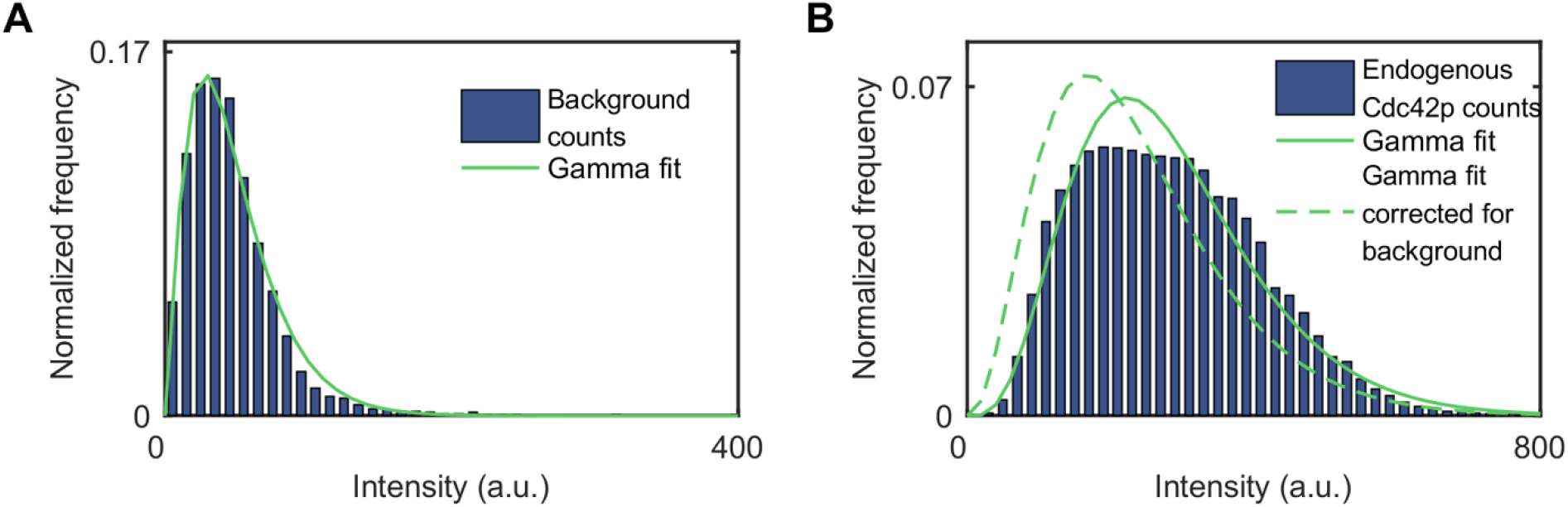
Measured Cdc42p distributions. (A) Flow cytometry data (histogram) of RWS116, which acts as a measure of the background count distribution without fluorescence. The solid green line shows the maximum likelihood gamma distribution fit. (B) Flow cytometry data (histogram) of RWS1421, which shows the WT distribution of Cdc42p. The solid green line shows the maximum likelihood gamma distribution fit, the dashed green line the fit corrected for the background fit in panel (A). Underlying data is found in Table S4.

**Table S1.**
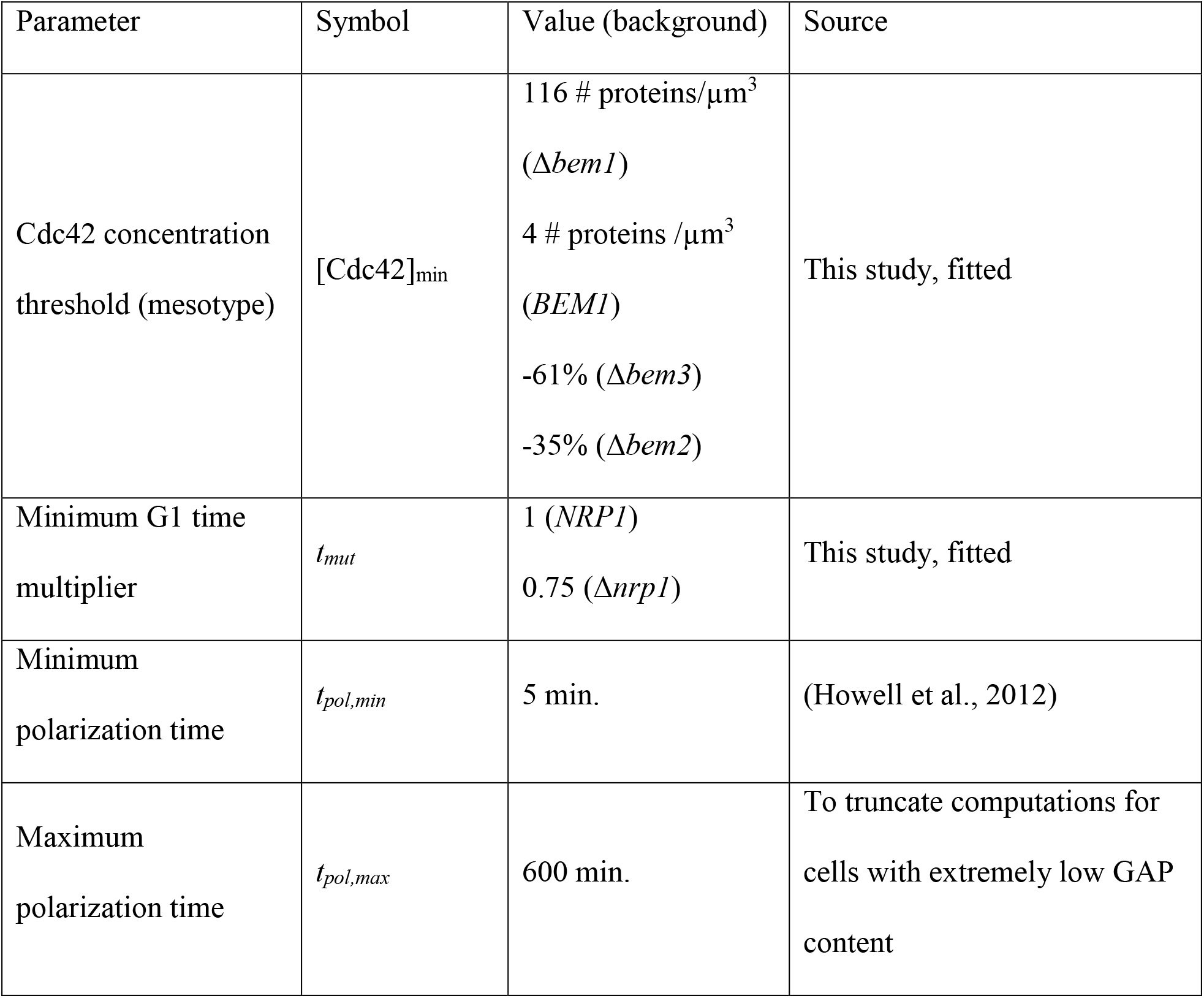
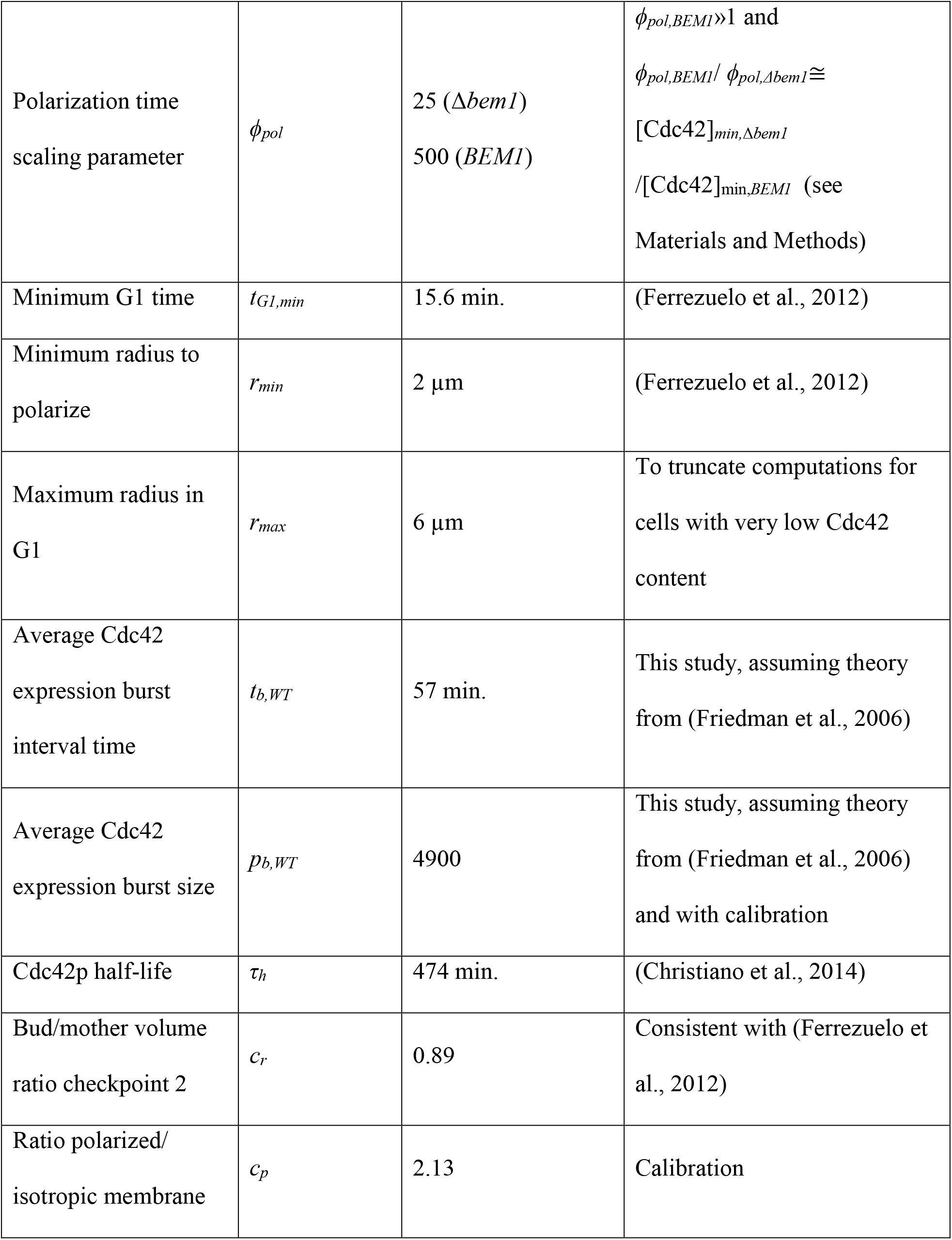
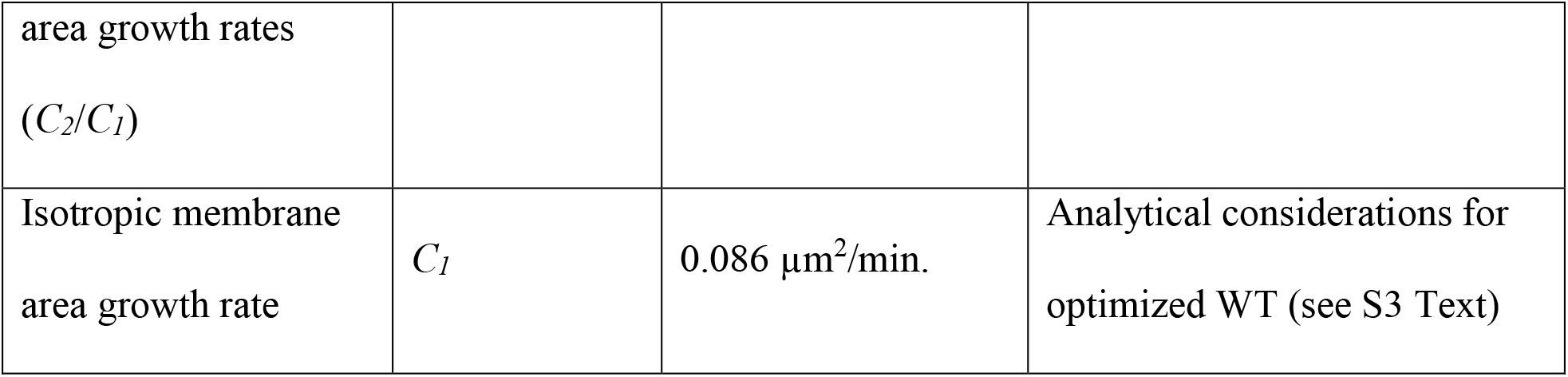
Growth model parameter list. Parameters used in the physical cell cycle model in this study, with associated symbols, values and justification.

**Table S2.** Additional G1 times shifts explained by stochastic minimum G1 time. Percentage of the mean shift in observed G1 times in the adaptive trajectory of (Laan et al., 2015) that is left unexplained by the standard cell cycle model implementation as in Figure 3D with minimum G1 time *t_mut_ t_G1,min_*, and is explained when the minimum G1 time is stochastic. In this alternative implementation, the minimum G1 time is on average the same, but results from a gamma distribution Γ(2, *t_mut_ t_G1,min_*/2).

| Genotype transition | % mean G1 time shift<br>mismatch described |
| --- | --- |
| WT $\rightarrow \Delta bem1$ | 5.8 |
| $\Delta bem1 \rightarrow \Delta bem1 \Delta bem3$ | 1.7 |
| $\Delta bem1 \Delta bem3 \rightarrow \Delta bem1 \Delta bem3 \Delta nrp1$ | 2.7 |
| $\Delta bem1 \Delta bem3 \Delta nrp1 \rightarrow$<br>$\Delta bem1 \Delta bem3 \Delta nrp1 \Delta bem2$ | 5.2 |

**Table S3.** Example calculation of quantitative epistasis of mutants with Δ*bem1*. Epistasis values for interactions with Δ*bem1* assuming several mutations with certain impact on model parameters. Epistasis is defined as relative multiplicative epistasis as in (da Silva et al., 2010). Default doubling time values are resulting from the fit in Figure 3A.

| Case | Parameter | Shift rel. to default [%] | Simulated doubling time [min.] | Simulated doubling time [min.] | Epistasis |
| --- | --- | --- | --- | --- | --- |
| Default | - | - | 82 | 406 | - |
| Fast membrane growth | $C_2$ | 10 | 77 | 460 | -0.08 |
| Slow membrane growth | $C_2$ | -10 | 89 | 368 | 0.07 |
| Small min. Start radius | $r_{min}, t_{G1,min}$ | -15,-20 | 70 | 234 | 0.17 |
| Large min. Start radius | $r_{min}, t_{G1,min}$ | 15,20 | 95 | 1018 | -0.34 |
| Slow in G1 | $t_{G1,min}$ | 20 | 94 | 1016 | -0.34 |
| Fast in G1 | $t_{G1,min}$ | -20 | 71 | 237 | 0.17 |
| Slow degradation | $\tau_h$ | 40 | 82 | 324 | 0.10 |
| Fast degradation | $\tau_h$ | -40 | 83 | 1038 | -0.41 |
| Small bursts | $p_{b,WT}$ | -5 | 82 | 555 | -0.14 |
| Large bursts | $p_{b,WT}$ | 5 | 82 | 327 | 0.09 |

**Table S4.** Flow cytometry data. Flow cytometry data underlying Fig S3 of RWS116 and RWS1421. The raw data, gated data and fitted data are in separate tabs. The fit of RWS1421 corrected for the background distribution fit of RWS116 is also given in the fitted data tab. The data in Table S1 underlies Figure S3.

**Table S5.** Growth assay data. Growth assay data of RWS1421 (OD_600_ values) with a weighted least squares fit on the log of the background corrected OD_600_ values, applied through our growth curve GUI.

**Table S6.** Growth assay data. Interaction data with *BEM1* from (Stark, 2006) and mutant data associating genes with various phenotypes and gene functions from (Cherry et al., 2012), SGD Project. https://www.yeastgenome.org/ [6 and 8 March 2018].

**Table S7.** Epistasis prevalence. Number of positive and negative interactions with *BEM1*, subdivided by mutant type.

|  | Number of negative<br>interactors with BEM1 | Number of positive<br>interactors with BEM1 |
| --- | --- | --- |
| Beneficial mutants (competitive<br>fitness) | 21 | 5 |
| Deleterious mutants (competitive<br>fitness) | 192 | 32 |
| Beneficial mutants (fermentative<br>growth) | 0 | 0 |
| Deleterious mutants (fermentative<br>growth) | 47 | 6 |
| Slow through G1 | 22 | 5 |
| Fast through G1 | 6 | 2 |
| Small at Start | 2 | 1 |
| Large at Start | 1 | 1 |
| Proteasomal | 5 | 2 |
| Ribosomal | 41 | 5 |
| Phospholipid | 5 | 2 |

**Table S8.** Oligonucleotides. Oligonucleotides used in this study.

**Data file S1. Flow cytometry processing code**

Matlab code to process flow cytometry data as found in Table S4.

**Data file S2. Yeast polarity GP map simulation code**

Zip-file containing the Matlab codes for the generation of the model simulation data.

**Data file S3. Plate reader GUI**

Zip-file containing the Matlab codes for analyzing growth assay data from excel sheets using a graphical user interface.

**Data file S4. Growth data effect of media quality**

Zip-file containing the excel sheets with measured OD_600_ values of the growth assays underlying Figure 4B.

