## Supplementary material for "A tractable bottom-up model of the yeast polarity genotype-phenotype map for evolutionary relevant predictions": Tables and code: Main_flow_cytometry_processing.m

```
function fitted_parameters = Main_flow_cytometry_processing()

% This function processes the flow cytometry data to calculate the Cdc42
% expression burst parameters for the paper entitled:
%
% "Tractable bottom-up modelling of the yeat polarity genotype-phenotype
% map to make evolutionary relevant predictions"
%
% Authors : Werner Karl-Gustav Daalman, Els Sweep, Liedewij Laan
%           Department of Bionanoscience, Faculty of Applied Sciences,
%           Delft University of Technology, Delft, the Netherlands
% Date    : 05-07-2021

%% Select raw data file interactively and load into variable 'Measurements'
[Data_filename,Data_pathname]   = uigetfile('.xlsx', 'Select flow cytometry raw data file (Table S4)');
% Pop-up menu to manually select file
if Data_filename == 0, return, end
% If nothing selected, quit
Data_filename_full              = [Data_pathname Data_filename];
Data_sheet                      = xlsread(Data_filename_full);
Background_data                 = Data_sheet(:, 1 : 3);
Endogenous_Cdc42_data           = Data_sheet(:, 4 : 6);
Background_data                 = Background_data(~isnan(Background_data(:, 1)), :);
Endogenous_Cdc42_data           = Endogenous_Cdc42_data(~isnan(Endogenous_Cdc42_data(:, 1)), :);

%% Gating (selecting the appropriate region in the Scattering plot (Forward vs. Side))
polygon         = [108541.266794626 33557.9514824798;  ....
                    103934.740882918 69946.0916442049; ...
                    237523.992322457 146765.498652291; ...
                    236948.176583493 91778.9757412399];

%% Apply polygon to filter out dead cells, junk, etc., yielding Gated_Data
ind                         = inpolygon(Background_data(:, 1), Background_data(:, 2), ...
                                polygon(:, 1),polygon(:, 2));
% Select data that is inside the polygon (gating)
Background_Gated_Data       = Background_data(ind, 3:end);
% Now remove glitch in some data points that state intensity of exactly 3, clearly artefact
Background_Gated_Data       = Background_Gated_Data(Background_Gated_Data ~= 3);

ind                         = inpolygon(Endogenous_Cdc42_data(:, 1),Endogenous_Cdc42_data(:, 2), ...
                                polygon(:,1),polygon(:,2)); %Logical array. Entry i equals
% Select data that is inside the polygon (gating)
Endogenous_Cdc42_Gated_Data = Endogenous_Cdc42_data(ind, 3:end);
% Now remove glitch in some data points that state intensity of exactly 3, clearly artefact
Endogenous_Cdc42_Gated_Data = Endogenous_Cdc42_Gated_Data(Endogenous_Cdc42_Gated_Data ~= 3);

% Note: this gated data is also present in the data file

%% Gamma fits (no deconvolution)
start_fit_background        = [2.4 20];
start_fit_endogenous        = [4.75 60];
% parameter starting points for fits

Background_Fits             = fitgamma(Background_Gated_Data, start_fit_background);
Endogenous_Fits             = fitgamma(Endogenous_Cdc42_Gated_Data, start_fit_endogenous);

%% Deconvolution of data with background (to remove background)
[k_s_endo, theta_s_endo]    = CalcparGam( Endogenous_Fits.k(2), Endogenous_Fits.theta(2), ...
                                          Background_Fits.k(2), Background_Fits.theta(2), ...
                                          Endogenous_Fits.k(4), Endogenous_Fits.theta(4), ...
                                          Background_Fits.k(4), Background_Fits.theta(4));
% element 2 in k/theta is the median value, element 4 is the error size in k/theta
fitted_parameters           = array2table([Background_Fits.k( 1: 3, 1)' ...
                                Background_Fits.theta( 1: 3, 1)'; ...
                                Endogenous_Fits.k( 1: 3, 1)' ...
                                Endogenous_Fits.theta( 1: 3, 1)'; ...
                                k_s_endo(:, 1)' theta_s_endo(:, 1)'], ...
                                'VariableNames', {'k_lower_bound', 'k', 'k_upper_bound', ...
                                'theta_lower_bound', 'theta', 'theta_upper_bound'}, ...
                                'RowNames', {'Background', 'Endogenous', ...
                                'Endogenous_minus_background'});

clearvars -except Background_Gated_Data Endogenous_Cdc42_Gated_Data fitted_parameters
save('Flow_cytometry_processed.mat');

end

function table_out = fitgamma(x, p0)

    % This function perofrms a maximum likelihood fit on the data in x
    % given starting parameters p0 ([k0, theta0]), and this fit is in the
    % output table_out, which also contains the 95% confidence interval

    pdf_gamma           = @(x, k1, theta1) gampdf(x, k1, theta1);
    % pdf of a gamma distribution
    options             = statset('MaxIter', 3000, 'MaxFunEvals', 6000);   
    lower_bounds        = [0 0];
    upper_bounds        = [Inf Inf];
    % bounds on k and theta
    [par_est, est_ci]   = mle(x, 'pdf', pdf_gamma, 'start', p0, 'lower', ...
                            lower_bounds, 'upper', upper_bounds, 'Options', options);
    var_out             = [est_ci(1, :); par_est; est_ci(2, :); (est_ci(2, :)-est_ci(1, :)) / 4];
    table_out           = table(var_out(:, 1), var_out(:, 2), 'VariableNames', ...
                            {'k' 'theta'}, 'RowNames', ...
                            {'Lower bound (95% CI)' ; 'Estimate'; 'Upper bound (95% CI)'; 'sigma'});
end

function [k_corr, theta_corr] = CalcparGam(k_endo, theta_endo, k_back, theta_back, ...
                                    s_k_endo, s_theta_endo, s_k_back, s_theta_back)

    % This function generates Monte Carlo samples of the background (gamma
    % distributed) corrected distribution of the endogenous gamma
    % distribution. This distribution can be approximated as a gamma
    % distribution by combining mean and variances, as long as the scaling
    % parameters theta are within an order of magnitude of each other [1].
    
    rng('default')
    
    mu_corrected    = @(k_e, t_e, k_b, t_b) k_e .* t_e - k_b .* t_b;
    sigma_corrected = @(k_e, t_e, k_b, t_b) k_e .* t_e.^2 - k_b .* t_b.^2;
    
    f_k_corr        = @(k_e, t_e, k_b, t_b) mu_corrected(k_e, t_e, k_b, t_b).^2 ./...
                                                sigma_corrected(k_e, t_e, k_b, t_b);
    f_theta_corr    = @(k_e, t_e, k_b, t_b) sigma_corrected(k_e, t_e, k_b, t_b) ./ ...
                                                mu_corrected(k_e, t_e, k_b, t_b);

    n               = 10000;

    % Sample draws for shape parameter k and scaling parameter theta,
    % assuming normal, independent errors on k and theta
    % Top draw is for zero noise on the fitted value
    sample_k        = f_k_corr( k_endo  + s_k_endo  .* [0; randn(n,1)], ...
                        theta_endo  + s_theta_endo  .* [0; randn(n,1)],...
                        k_back  + s_k_back  .* [0; randn(n,1)], ...
                        theta_back  + s_theta_back  .* [0; randn(n,1)]);
    sample_theta    = f_theta_corr( k_endo  + s_k_endo  .* [0; randn(n,1)], ...
                        theta_endo  + s_theta_endo  .* [0; randn(n,1)], ...
                        k_back  + s_k_back  .* [0; randn(n,1)], ...
                        theta_back  + s_theta_back  .* [0; randn(n,1)]);
    
    k_corr          = [quantile(sample_k(2:end), 0.025); sample_k(1); ...
                        quantile(sample_k(2:end), 0.975)];
    theta_corr      = [quantile(sample_theta(2:end), 0.025); sample_theta(1); ...
                        quantile(sample_theta(2:end), 0.975)];

end

%% Bibliography
%[1]    Stewart T, Strijbosch LWG, Moors H, Batenburg P van. A simple
%       approximation to the convolution of gamma distributions. 2007;
```
